# OGDHL regulates tumor growth, neuroendocrine marker expression, and nucleotide abundance in prostate cancer

**DOI:** 10.1101/2025.05.28.656673

**Authors:** Matthew J. Bernard, Angel Ruiz, Johnny A. Diaz, Nicholas M. Nunley, Rachel N. Dove, Shile Zhang, Ernie Lee, Kylie Y. Heering, Sachi Bopardikar, Andrea Gallardo, Takao Hashimoto, Raag Agrawal, Chad M. Smith, Blake R. Wilde, Nedas Matulionis, Helen M. Richards, Sandy Che-Eun S. Lee, Marina N. Sharifi, Joshua M. Lang, Shuang G. Zhao, Michael C. Haffner, David B. Shackelford, Paul C. Boutros, Heather R. Christofk, Andrew S. Goldstein

## Abstract

As cancer cells evade therapeutic pressure and adopt alternate lineage identities not commonly observed in the tissue of origin, they likely adopt alternate metabolic programs to support their evolving demands. Targeting these alternative metabolic programs in distinct molecular subtypes of aggressive prostate cancer may lead to new therapeutic approaches to combat treatment-resistance. We identify the poorly studied metabolic enzyme Oxoglutarate Dehydrogenase-Like (OGDHL), named for its structural similarity to the tricarboxylic acid (TCA) cycle enzyme Oxoglutarate Dehydrogenase (OGDH), as an unexpected regulator of tumor growth, treatment-induced lineage plasticity, and DNA Damage in prostate cancer. While OGDHL has been described as a tumor-suppressor in various cancers, we find that its loss impairs prostate cancer cell proliferation and tumor formation. Loss of OGDHL profoundly alters Androgen Receptor inhibition-induced plasticity, including suppressing the neuroendocrine markers DLL3 and HES6, induces accumulation of the DNA damage response marker ƔH2AX, and reduces nucleotide synthesis. Our data suggest that OGDHL has minimal impact on TCA cycle activity, and that mitochondrial localization is not required for its regulation of prostate cancer plasticity and nucleotide metabolism. Finally, we demonstrate that OGDHL expression is tightly correlated with neuroendocrine differentiation in clinical prostate cancer. These findings underscore the importance of investigating poorly characterized metabolic genes as potential regulators of distinct molecular subtypes of aggressive cancer.

## Introduction

Cells constantly undergo rapid metabolic changes to respond to external stimuli or local nutrient availability, and to meet biosynthetic demands^1^. This ability to quickly overhaul metabolic networks allows healthy cells to properly integrate signaling, regulate redox homeostasis, and modulate epigenetic substrates that dictate cell fate decisions^2,3^. Due to differences in bioenergetic and metabolite pool needs, these networks are fine tuned in a cell-type specific manner. During oncogenic transformation and disease progression, cancer cells hijack cellular metabolism to evade therapeutic and biological pressures^4,5^. There is a growing appreciation for the reciprocal relationship between cellular lineage and metabolism^6–8^. Perturbation of these metabolic switches dramatically alters cell identity and differentiation. One major function of metabolic enzymes is to generate energy and fuel anabolic growth. In addition, recent findings have indicated that many of these same enzymes also serve non-canonical functions that more directly regulate transcriptional profiles and cellular signaling^9–12^.

Prostate cancer is one of the most common cancers globally, accounting for nearly 1.5 million new cases annually^13^. In the United States alone, there will be more than 35,750 deaths in 2025^14^. Identifying novel strategies for the treatment of advanced prostate cancer is essential for improving outcomes for patients with prostate cancer. Because Androgen Receptor (AR) signaling can promote survival and proliferation of tumor cells^15^, most advanced prostate cancers are treated with hormonal therapies that target the AR signaling axis, including Enzalutamide and Apalutamide that directly bind and inhibit AR. While AR inhibitors have improved patient survival for individuals with CRPC^16,17^, disease recurrence is nearly universal. One mechanism through which tumor cells adapt to AR blockade is through changes in lineage identity that result in AR-indifferent tumors^18^. Despite recent promising clinical trial results^19^, tumors that become resistant to AR blockade currently lack many effective therapeutic options and are almost uniformly lethal^20^. Understanding how cells adapt to AR inhibition and drive resistant phenotypes is essential for developing new therapies for patients with advanced treatment-resistant prostate cancer.

We previously demonstrated that castration-resistant prostate cancer (CRPC) cells undergo extensive metabolic remodeling in response to prolonged AR inhibition using the antiandrogen drug Enzalutamide^21^. Notably, cells adapting to sustained AR blockade exhibit altered mitochondrial morphology and increased reliance on mitochondrial oxidative metabolism. In prostate cancer, metabolic rewiring occurs concomitantly with the activation of AR-independent lineage programs to evade pharmacological targeting, including acquisition of neuronal, neuroendocrine (NE), and stem-like features^22–26^. We know little about whether treatment-induced changes in metabolism directly regulate treatment-induced plasticity phenotypes, and which enzymes may be involved in this process. The mitochondria serve as a central hub for both bioenergetic production and metabolite pool regeneration, while also influencing cell signaling^27^. Disruption of metabolite pools can induce apoptosis, alter cell fate, and exacerbate cancer progression through the accumulation of oncometabolites. One metabolic enzyme associated with mitochondrial function is Oxoglutarate Dehydrogenase-Like (OGDHL), an isozyme of Oxoglutarate Dehydrogenase (OGDH), which catalyzes the rate-limiting step in the interconversion of alpha-ketoglutarate (a-KG) into succinyl-CoA within the tricarboxylic acid (TCA) cycle. While OGDH is expressed robustly throughout nearly every cellular lineage in the body, OGDHL expression is primarily localized to the brain and liver^28,29^. OGDHL is expressed in various cancer types and displays tumor suppressor properties in kidney, cervical, and pancreatic cancers^30–35^. Despite its aberrant expression across these multiple cancers, our understanding of the function of OGDHL remains poorly characterized. In a few studies, OGDHL has been suggested to modulate immune signaling, lipid metabolism, and DNA damage^30,34,35^.

Replication of DNA for proliferation is a strictly regulated process due to the deleterious nature of most mutations. High fidelity replication of nearly 3 billion base pairs is required each time a cell proliferates. Because of this, the DNA repair pathway is critical to maintain cell health and proper proliferation. Many of the most commonly altered tumor suppressor genes, including *Tp53*, *Brca2*, and *Rb1*, are regulators of the DNA damage response and/or serve as cell cycle checkpoints^36–38^. Mutations in these critical genes can usurp the DNA damage regulatory mechanisms and unleash unchecked cell growth, resulting in a more frequent mutation rate and increased susceptibility to oncogenic transformation^39^. Cells accumulate the metabolite precursors required for DNA synthesis through several mechanisms including salvage of ribonucleotides and deoxyribonucleotides from the surrounding microenvironment and through *de novo* synthesis pathways^40^. Rewiring of these metabolic pathways through alternative means during biological and therapeutic pressures serves as one way tumor cells adapt to sustain tumor growth^41^. Precise regulation of these nucleotide pools is paramount for proper RNA and DNA synthesis and repair pathways. Imbalance of these pools can induce replication stress, leading to cell cycle arrest and apoptosis. Dysregulation of nucleotide pools has also been linked to altered lineage differentiation, which is associated with sensitivity or resistance to targeted therapies. In summary, the intersection of metabolic rewiring and nucleotide metabolism represents an intriguing target for the treatment of aggressive and therapy-resistant cancer subtypes.

Here, we identify the TCA cycle enzyme OGDHL as a regulator of tumor cell proliferation and treatment-induced lineage plasticity in prostate cancer. OGDHL is elevated in a group of prostate cancer cell lines, patient-derived xenograft (PDX) models, genetically engineered mouse models, and clinical tissues from patients with aggressive, lethal treatment-resistant prostate cancer. Unlike in prior reports of other cancer-types, we did not find evidence for OGDHL as a tumor suppressor or as a regulator of the TCA cycle. Instead, loss of OGDHL expression impairs cell proliferation and tumor formation. We demonstrate that chronic loss of OGDHL restrains the treatment-associated plasticity in response to prolonged AR pathway inhibition, reducing expression of NE markers HES6 and DLL3. Loss of OGDHL induces accumulation of the DNA damage marker ƔH2AX and reduces nucleotide synthesis, resulting in depleted nucleotide pools. Mechanistically, we found that OGDHL function does not require mitochondrial localization to maintain its role in regulation of lineage marker expression and nucleotide metabolism in prostate cancer.

## Results

### OGDHL regulates cell proliferation and tumor growth in castration-resistant prostate cancer

Having established that prolonged AR inhibition induces a metabolic shift characterized by increased reliance on mitochondrial oxidative metabolism *in vitro*^21^, we evaluated the effect of AR blockade on tumor cell metabolism *in vivo* using a multi-tracer Positron Emission Tomography (PET) imaging strategy. We generated xenograft tumors from MDA-PCa 180-30^42^ cells in NOD-SCID-IL2Rg^null^ (NSG) mice and treated with enzalutamide via daily oral gavage for 2 weeks. To define treatment-induced changes in tumor metabolism *in vivo*, we first conducted radiolabeled tracing using ^18^F-FBnTP-PET to measure mitochondrial membrane potential as an indicator of oxidative capacity. Twenty-four hours later, we performed ^18^F-FDG-PET to measure glucose uptake as a surrogate for glycolytic activity. Enzalutamide treatment induced a statistically significant reduction in ^18^F-FDG signal (**Figure 1A**), while ^18^F-FBnTP signal was not significantly different (**Figure 1B**), indicating that enzalutamide treatment also induces a metabolic shift toward mitochondrial oxidative metabolism *in vivo*. Two weeks of Enzalutamide treatment in MDA-PCa 180-30 tumor-bearing mice was sufficient to reduce tumor size and weight (**Figure S1, S1B)**. As we determined that increased reliance on mitochondrial oxidative metabolism was a consequence of AR blockade *in vitro* and *in vivo*, we evaluated transcriptional changes in TCA cycle enzymes following extended (> 6 weeks) Enzalutamide treatment. OGDHL was the most enriched enzyme, exhibiting a greater than 2-fold increase in mRNA expression in Enzalutamide-maintained cells compared to control 16D^CRPC^ cells (**Figure 1C**). We next verified that OGDHL protein expression was elevated in response to Enzalutamide treatment and found that this upregulation of OGDHL did not correspond to a compensatory downregulation of the homologous OGDH (**Figure 1D**). OGDHL protein expression was roughly 30% greater in Enzalutamide-treated MDA-PCa 180-30 CRPC tumors relative to vehicle-treated control tumors (**Figure 1E, 1F)**.

**Figure 1:**
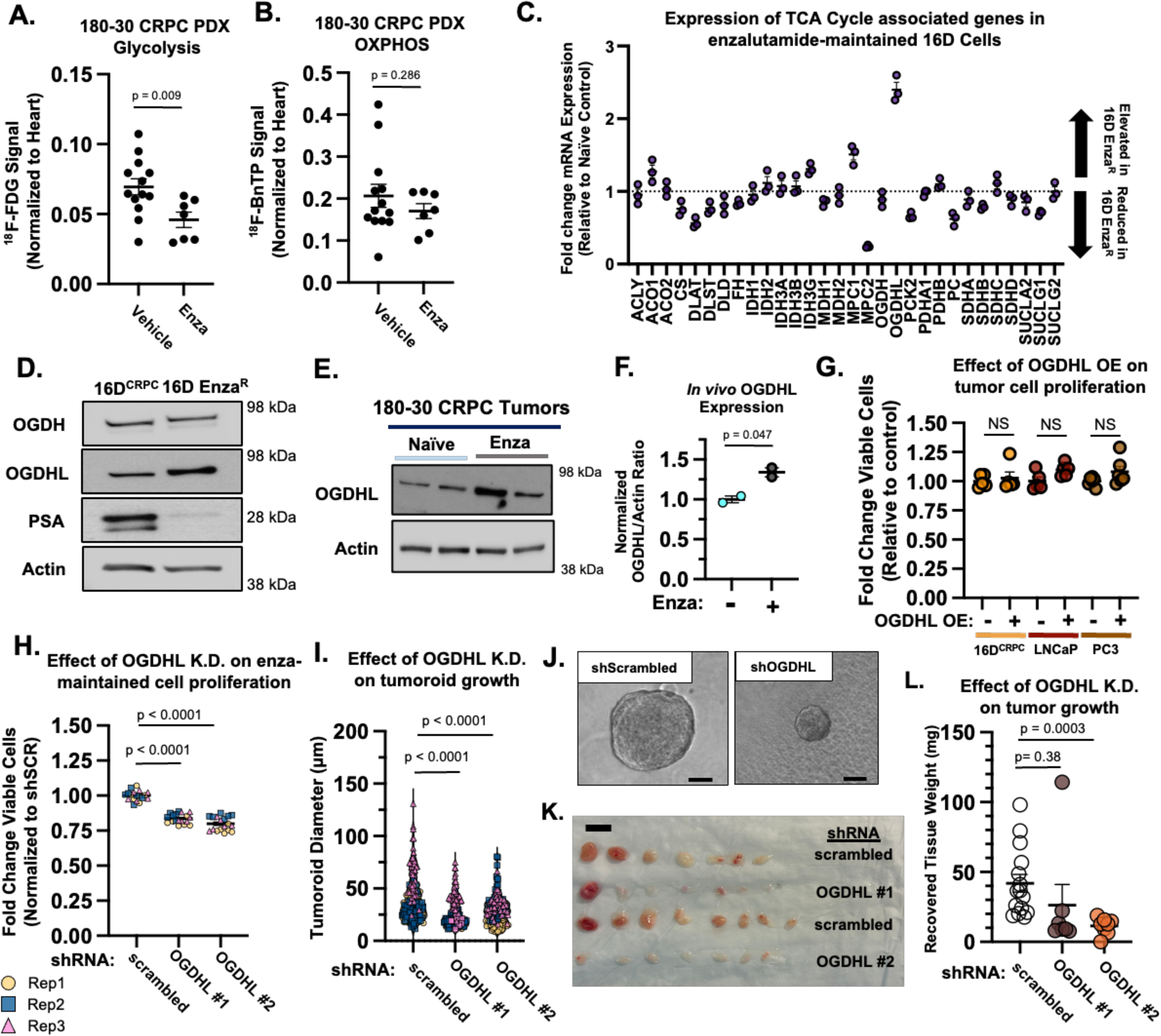
OGDHL supports proliferation of Enzalutamide-resistant prostate cancer cells *in vitro* and *in vivo*. **(A and B)** ^18^F-FDG PET signal **(A)** or ^18^F-BnTP PET signal **(B)** in MDA PCa 180-30 CRPC tumors treated with Enza or Vehicle for 14 Days. **(C)** RNA expression of Canonical TCA cycle genes in Enza-Resistant (Enza^R^) 16D Cells relative to Enza-naïve cells. **(D)** Western Blot of changes in OGDHL, PSA and OGDH in response to acquisition of Enza-resistance in 16D^CRPC^ cells. **(E and F)** Western Blot of elevated OGDHL expression in MDA PCa 180-30 derived xenografts **(E)**, and quantification of *in vivo* western blot **(F)**. **(G)** Relative change in cell viability over 48 hours with OGDHL overexpression. Plotted as change in viability relative to control cells. Data from 5 (16D, LNCaP) or 6 (PC3) technical replicates. **(H)** Relative change in cell viability over 48 hours with shRNA-mediated knockdown of OGDHL in Enza-Maintained 16D cells. Plotted as change in viability relative to control. Data from 3 biological replicate experiments with 5 technical replicates each. **(I)** Measured tumoroid diameter of Enza-maintained 16D cells after 2 weeks of 3D culture. Data from 3 biological replicate experiments, >50 tumoroids measured per replicate. **(J)** Representative images of control (shScrambled) and OGDHL knockdown (shOGDHL) tumoroids generated from Enza-Maintained 16D Cells. Scale Bar = 50 μm. **(K and L)** Image of tissues recovered 4 weeks after implantation *in vivo* from Enza-maintained 16D^CRPC^ cells with control (Scrambled) or OGDHL knockdown. Scale Bar = 1.0 cm. **(K)** Measured weights of tissues recovered 4 weeks after implantation *in vivo* from Enza-maintained 16D Cells with control (Scrambled) or OGDHL knockdown **(L)**. Error bars represent +/- SEM. P-value calculated by unpaired t-test with Welch’s Correction.

Although the precise physiological role of OGDHL in the TCA cycle has not been well defined, OGDHL has been investigated as a prognostic biomarker for multiple cancers^43,44^ and is reported to have tumor-suppressive activity in cervical, pancreatic, kidney, and liver cells^30–35^. To investigate the role of OGDHL in prostate cancer, we maintained three parallel sub-lines of 16D^CRPC^ cells in Enzalutamide to allow them to independently adapt to prolonged AR inhibition. OGDHL protein levels increased in each of these lines and this expression increased over time as cells adapted to AR blockade (**Figure S1C**). OGDHL expression was also increased in response to AR inhibition in two independent *in vitro* models of CRPC: LuCAP35^CR^ cells grown in 2D culture, and MDA-PCa 180-30^42^ organoids (**Figure S1D, S1E)**. To determine the functional role of OGDHL in prostate cancer, we generated constitutive overexpression systems in several prostate cancer cell line models (**Figure S1F**). Forced expression of OGDHL had no impact on tumor cell viability in LNCaP, 16D^CRPC^ or PC3 cells (**Figure 1G**). Because increased OGDHL expression seemed to be a consistent cellular response to prolonged AR blockade, we next evaluated how loss of OGDHL influences proliferation. Short hairpin RNA (shRNA)-mediated knockdown of OGDHL in Enzalutamide-maintained 16D^CRPC^ and LuCAP35^CR^ cells significantly reduced cell viability (**Figure 1H, S1G-I**). We implanted control and knockdown cells in Matrigel to generate 3D tumoroids and found that OGDHL knockdown significantly reduced tumoroid size, an indicator of proliferative capacity (**Figure 1I, 1J**).

To evaluate the effect of OGDHL loss on tumor formation, we implanted control and OGDHL knockdown Enzalutamide-maintained 16D^CRPC^ cells with Matrigel into NSG mice and quantified outgrowths after 4 weeks *in vivo*. Tissues recovered from implantation of OGDHL knockdown cells were significantly smaller. Recovered tissues had an average mass of 26.26 +/- 36.24 mg (shOGDHL #1) and 11.34 +/- 5.57 mg (shOGDHL #2) compared to 41.86 +/- 23.67 mg in scrambled control tissues (**Figure 1K, 1L).** Most of the tissues derived from knockdown cells lacked visible outgrowths, so we quantified the percentage of recovered tissues weighing more than the Matrigel alone. Only 33.3% (5/15) of tissues resected from OGDHL knockdown xenografts weighed more than the injection medium alone, compared to 100% (15/15) of tissues resected from control cells (p = 0.0002), suggesting that OGDHL knockdown cells fail to efficiently grow *in vivo*. We validated reduced protein abundance of OGDHL in the recovered tissues from knockdown cells (**Figure S1J**). Interestingly, one outgrowth from OGDHL knockdown cells was greater than five times heavier than any other OGDHL knockdown tissue, suggesting that in rare instances, cells can overcome OGDHL loss to form tumors *in vivo*. Taken together, our data suggest that OGDHL does not act as a tumor-suppressor in CRPC, as has been shown in other cancer contexts. These findings indicate that OGDHL supports CRPC growth *in vitro* and tumor-forming capacity *in vivo*.

### OGDHL loss suppresses cell cycle-related signatures in vitro and in vivo

Because the role of OGDHL in CRPC seems to differ from its reported function in other cancers, we investigated the effect of OGDHL loss on gene expression by conducting RNA-sequencing in Enzalutamide-Maintained 16D^CRPC^ cells. We first asked whether OGDHL loss induces compensatory changes in the expression of closely related TCA cycle enzymes. We found minimal changes in expression of canonical TCA cycle genes (**Figure 2A**) or a broader list of TCA cycle-associated genes (**Figure S2A**) in response to acute OGDHL loss. We next took an unbiased approach to determine broad transcriptional changes induced by acute knockdown of OGDHL. Significantly upregulated genes were associated with cell stress pathways, including *GADD45A*, *CASP4*, and *TP53INP2*. Significantly downregulated genes were associated with cell cycle and proliferation, such as *CDC20*, *CENPN*, *MYC*, and *TOP2A* (**Figure 2B, 2C**). Gene Set Enrichment Analysis (GSEA) revealed strong downregulation of cell cycle signatures, including the Hallmark gene sets for G2M checkpoint, E2F targets, and Myc Targets at both the gene set and individual gene level (**Figures 2E, S2B**). To confirm results obtained using shRNA-mediated knockdown, we used a CRISPR-Cas9 approach to generate independent OGDHL knockout 16D^CRPC^ cell lines that are maintained in Enzalutamide. After validating the knockout cells by Western blot (**Figure S2C**), we found that chronic loss of OGDHL did not induce compensatory upregulation of TCA cycle genes (**Figures S2D, S2E**). Similar to acute knockdown of OGDHL, genetic knockout of OGDHL led to a pronounced reduction in G2M checkpoint genes, E2F target genes, and Myc target genes (**Figure 2E**). We implanted OGDHL knockout and control cells in castrated NSG mice to maintain suppression of AR signaling *in vivo* and performed RNA sequencing upon tumor formation. After validating loss of OGDHL protein (**Figure S2F**), we found a strong transcriptional downregulation of G2M checkpoint, E2F targets, and Myc Target genes in knockout cells *in vivo* (**Figure 2F**). Taken together, these data indicate that loss of OGDHL in AR inhibition-resistant CRPC cells represses transcription of cell cycle associated genes and activates transcriptional pathways related to cellular stress.

**Figure 2:**
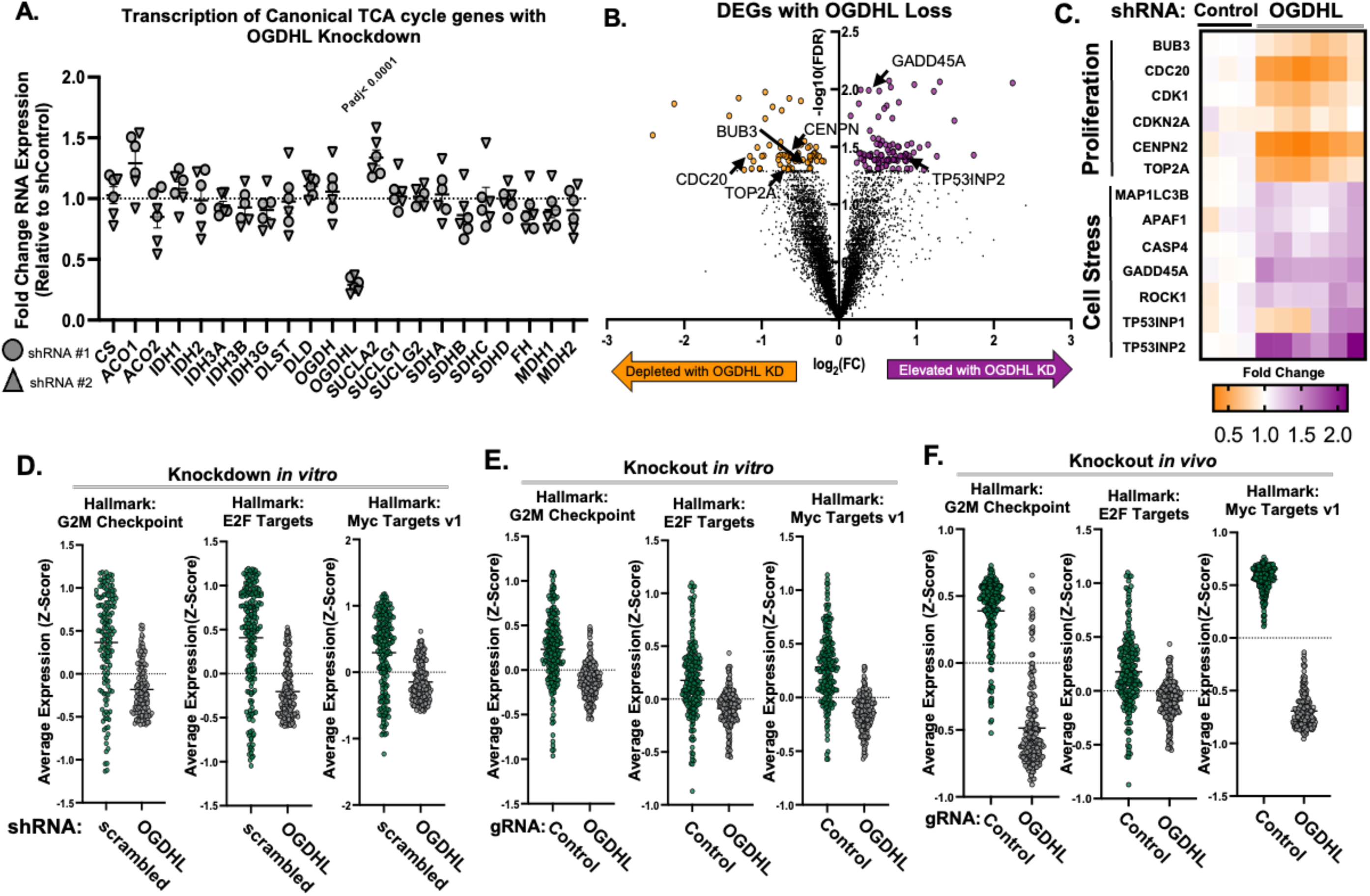
Loss of OGDHL in CRPC cells alters transcription of cell stress and cell cycle associated pathways *in vitro* and *in vivo*. **(A)** RNA expression of Canonical TCA cycle genes in OGDHL knockdown Enza-maintained 16D^CRPC^ cells *in vitro* normalized to control knockdown values. Data represented as 3 technical replicates each from 2 shRNAs. **(B)** Volcano Plot of Differentially Expressed Genes in shOGDHL Enza-maintained 16D^CRPC^ cells relative to control. Color indicates FDR < 0.05 **(C)** Heatmap of isolated proliferation and cell stress genes in control and OGDHL knockdown cells, with technical replicates shown. **(D-F)** Average expression of genes in Hallmark: G2M Checkpoint, E2F Targets, and Myc Targets v1 with Enza-Maintained 16D^CRPC^ cells with acute knockdown of OGDHL *in vitro* **(D)**, in OGDHL CRISPR knockout cell lines *in vitro* **(E)**, and in tumors formed from OGDHL knockout cells **(F)**. Data shown is average Z scores from 3 control replicates and 6 knockdown/knockout replicates *in vitro* or 5 control and 4 knockout replicates *in vivo.* Error bars represent +/- SEM. Adjusted P values were calculated by applying the Benjamini-Hochberg procedure for multiple hypothesis testing to p-values corresponding to 2-tailed t-tests.

### OGDHL regulates nucleotide biosynthesis in Enzalutamide-treated castration-resistant prostate cancer

Due to the unexpected effect of OGDHL loss on transcription of proliferation and cellular stress genes, we next set out to determine the metabolic function of OGDHL in prostate cancer cells. OGDHL is predicted to form a complex with dihydrolipoyl succinlyltransferase (DLST) and dihydrolipoyl dehydrogenase (DLD) to catalyze the conversion of alpha-ketoglutarate (a-KG) into succinyl-CoA in the TCA cycle^45^. To measure changes in nutrient utilization, we conducted heavy-isotope nutrient tracing experiments using U-^13^C Glucose and U-^13^C Glutamine for 24 hours (**Figure 3A**) in Enzalutamide-Maintained 16D^CRPC^ cells following OGDHL knockdown. OGDHL loss did not lead to an accumulation of heavy-isotope labeled a-KG (**Figure 3B**) or a reduction in labeled carbons downstream of a-KG from either glucose or glutamine (**Figure S3A-S3C**). These findings suggest that OGDHL plays a limited role in regulating the incorporation of glucose or glutamine-derived metabolites into the TCA cycle in CRPC cells, possibly due to the functional redundancy with OGDH.

**Figure 3:**
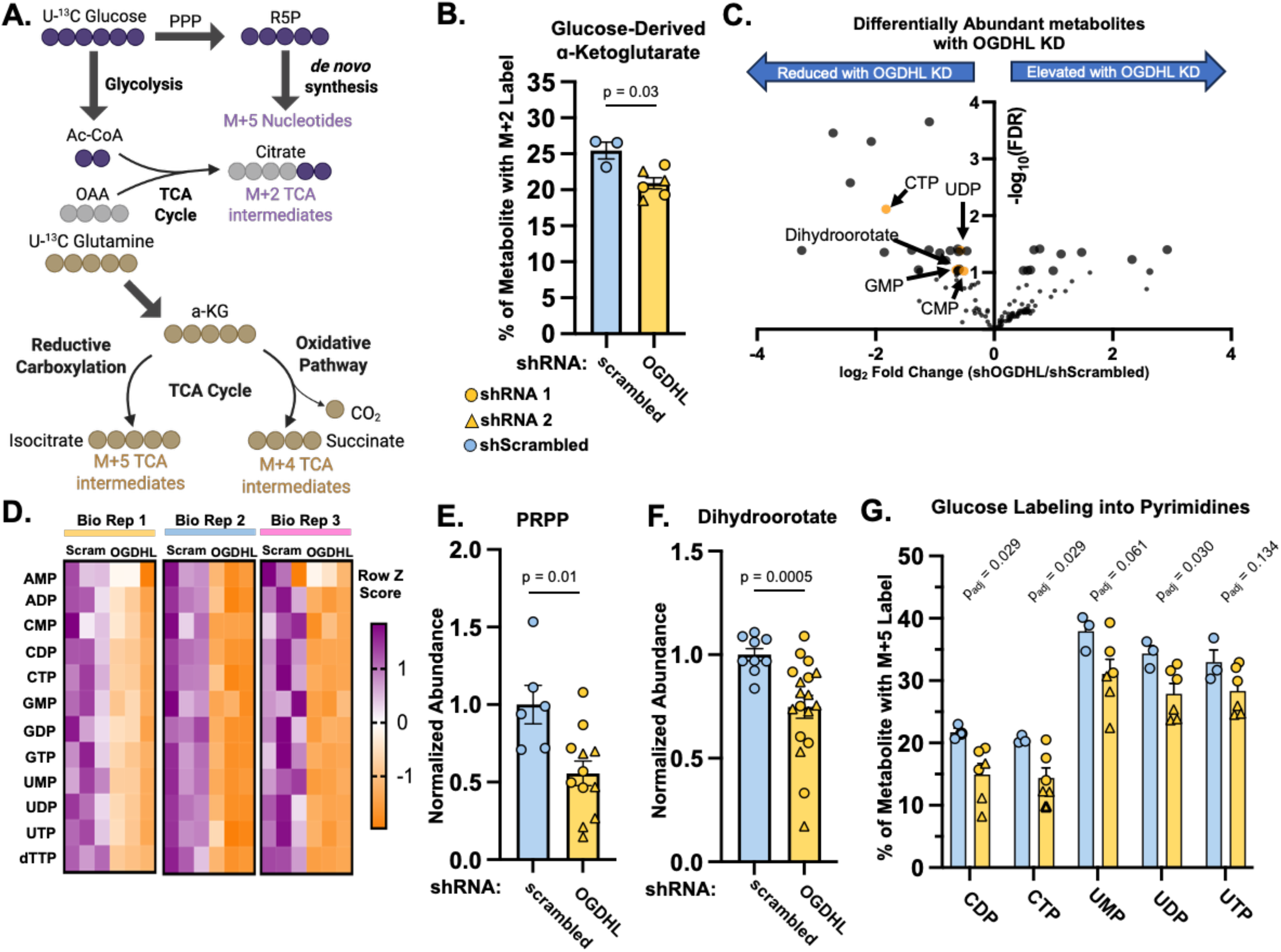
OGDHL loss perturbs nucleotide pools and glucose incorporation into nucleotides. **(A)** Schematic of heavy-isotope (U-^13^C Glucose or U-^13^C Glutamine) nutrient tracing into TCA cycle and nucleotide metabolic intermediates. **(B)** M + 2 labeled ɑ-ketoglutarate from U-^13^C glucose in control or OGDHL knockdown Enza-maintained 16D^CRPC^ cells. Data represent 3 technical replicate experiments for each of 2 shRNAs. **(C)** Volcano Plot of differentially abundant metabolites from metabolic profiling of Enza-maintained 16D^CRPC^ cells with OGDHL or control knockdown. Highlighted metabolites are nucleotide synthesis intermediates. Line indicates FDR = 0.05. **(D)** Heatmap of nucleotide phosphate abundance in OGDHL knockdown and control knockdown Enza-maintained 16D^CRPC^ cells. Data represented as row z scores from 3 technical replicates from each of 3 biological replicate lines. **(E and F)** Relative abundance of nucleotide synthesis intermediates Phosphoribosyl pyrophosphate (PRPP) **(E)** and Dihydroorotate **(F)** from metabolic profiling in control and OGDHL knockdown Enza-maintained 16D^CRPC^ cells. Data from 2 biological replicate experiments (PRPP) or 3 biological replicates (Dihydroorotate) each with 3 technical replicates. **(G)** M + 5 labeling of pyrimidine phosphates derived from U-^13^C glucose in control and OGDHL knockdown Enza-maintained 16D^CRPC^ cells. Error bars represent +/- SEM. Adjusted P values were calculated by applying the Benjamini-Hochberg procedure for multiple hypothesis testing to p-values corresponding to 2-tailed t-tests.

We next attempted to identify other metabolic consequences of acute OGDHL loss. Many of the significantly depleted metabolites were involved in pyrimidine biosynthesis and nucleotide homeostasis (**Figure 3C**). Across multiple biological replicates of Enzalutamide-maintained prostate cancer cells, OGDHL loss consistently led to a depletion of nucleotide phosphates (**Figure 3D**) and reduced abundance of certain biosynthetic intermediates of nucleotide synthesis including Phosphoribosyl pyrophosphate (PRPP) and dihydroorotate **(Figure 3E, 3F**). Glucose can be incorporated into *de novo* synthesized nucleotides through conversion to a ribose backbone via the pentose phosphate pathway (**Figure 3A**). We found that acute OGDHL loss reduced carbon labeling from glucose into newly synthesized nucleotides phosphates, including both pyrimidines **(Figure 3G**) and purines **(Figure S3D).** This reduction in labeling was concurrent with a reduction in nucleotide pools, consistent with decreased activity of the pentose phosphate pathway and diminished *de novo* nucleotide synthesis.

To understand how chronic loss of OGDHL influences cellular metabolism, we conducted heavy-isotope nutrient tracing experiments in OGDHL knockout cell lines. When we tracked glucose incorporation into the TCA cycle, we found that labeling of a-KG was similar to labeling of other TCA cycle intermediates (**Figure S3E).** Further, OGDHL knockout did not alter labeling from glutamine into the TCA cycle, through either the oxidative (**Figure S3F**) or reductive carboxylation (**Figure S3G)** pathways. Much like in response to acute knockdown, metabolic profiling of OGDHL knockout cells revealed a reduction in abundance of ribonucleotides and deoxyribonucleotides (**Figure S3H**). Taken together, our metabolomic data indicate that the main metabolic consequence of OGDHL loss in CRPC cells is not altered TCA cycle metabolism, but rather depletion of nucleotide pools, at least in part through slowed *de novo* nucleotide synthesis.

### OGDHL regulates DNA repair and cell stress pathways in prostate cancer

Nucleotide imbalance has a profound impact on cellular proliferation, cell cycle progression, and DNA damage machinery^46^. As we found that loss of OGDHL diminished nucleotide pools and reduced glucose incorporation into newly synthesized nucleotides, we evaluated expression of nucleotide metabolism genes. Genetic knockout of OGDHL *in vitro* and *in vivo* reduced expression of critical *de novo* nucleotide synthesis and salvage genes including Carboamoyl-Phosphate Synthase 2 (*CAD*), Dihydroorotate Dehydrogenase (*DHODH*), Uridine monophosphate synthetase (*UMPS*), Hypoxanthine phosphoribosyltransferase 1 (*HPRT1)* and Adenine phosphoribosyl transferase (APRT) (**Figure 4A, 4B, S4A, S4B**). More broadly, OGDHL knockout reduced expression of genes in the KEGG Pyrimidine Metabolism gene set (**Figure 4C**). Similarly, we found that acute knockdown of OGDHL in Enzalutamide-maintained 16D^CRPC^ cells reduced transcription of *de novo* pyrimidine synthesis and purine salvage genes (**Figure S4C)**. These data indicate that diminished nucleotide abundance following the loss of OGDHL expression in CRPC is likely caused by impaired *de novo* synthesis and nucleotide salvage.

**Figure 4:**
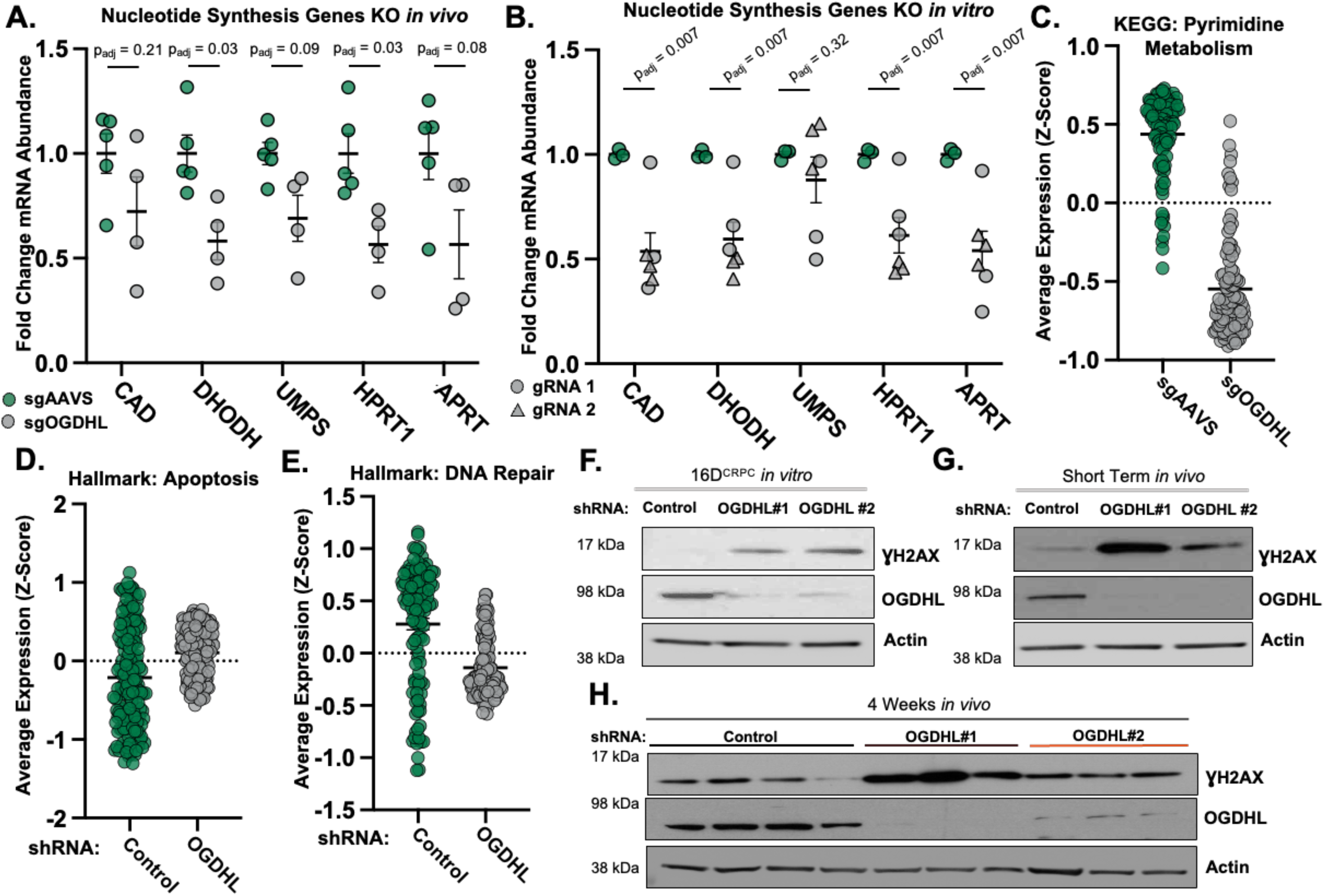
OGDHL Loss reduces nucleotide synthesis gene expression and induces DNA damage *in vitro* and *in vivo*. **(A and B)** Relative RNA abundance of nucleotide synthesis genes in tumors formed from 16D^CRPC^ cells with genetic knockout of OGDHL (n =4) or a control (n = 5) **(A)** or OGDHL knockout cells grown *in vitro* **(B).** Data from 3 technical replicates each from 2 gRNAs. **(C)** Average Expression of genes in the KEGG: Pyrimidine metabolism gene set in 16D^CRPC^ cells with genetic knockout of OGDHL. Data shown is average Z scores from 3 control replicates and 6 knockout replicates *in vitro.* **(D and E)** Average expression of genes in the Hallmark: Apoptosis **(D)** and Hallmark: DNA Repair **(E)** gene sets with acute knockdown of OGDHL. Data shown is average Z scores from 3 control replicates and 6 knockdown replicates *in vitro.* **(F-H)** Western blot of the DNA damage marker ƔH2AX and OGDHL expression in control and OGDHL knockdown cells *in vitro* **(F)**, in cells recovered 4 days after implantation *in vivo* **(G)** and in cells recovered 4 weeks after implantation *in vivo* **(H).** Error bars represent +/- SEM. Adjusted P values were calculated by applying the Benjamini-Hochberg procedure for multiple hypothesis testing to p-values corresponding to 2-tailed t-tests.

Depletion of nucleotides can cause genomic instability and trigger replication stress, leading to double stranded DNA breaks, cell cycle arrest and apoptosis^47,48^. OGDHL has been implicated as a potential regulator of DNA damage in cancer^35,49^. Thus, we wondered whether loss of OGDHL may cause increased replication stress. Acute knockdown of OGDHL led to increased transcription of genes in the Hallmark Apoptosis gene set and reduced expression of DNA repair pathway genes (**Figure 4D, 4E).** In addition to transcriptional hallmarks of replication stress, we also identified that knockdown of OGDHL *in vitro* led to increased abundance of ƔH2AX (**Figure 4F**), a sensitive marker of double stranded DNA breaks and a hallmark of DNA damage^50^. We similarly observed an increase in ƔH2AX when OGDHL knockdown cells were grown *in vivo* (**Figure 4G, 4H, S4E**). We confirmed that OGDHL knockdown induces DNA damage using a comet assay based on single cell gel electrophoresis (**Figure S4F**). OGDHL loss also reduced expression of the DNA repair pathway genes CHK1 and CHK2 (**Figure S4G).** As DNA damage can induce cell death, we questioned whether OGDHL loss induced higher rates of apoptosis. We found that OGDHL loss increased the percentage of Annexin V^+^ cells (**Figure S4H)**. To confirm that these phenotypes are driven by OGDHL loss, we introduced a knockdown-resistant version of OGDHL and found that the rescue reduced markers of DNA damage including DNA tail length by the comet assay and ƔH2AX (**Figure S4F, S4I**). We wondered whether OGDHL-loss induced DNA damage accumulation and apoptosis could be related to depletion of nucleotides. Supplementation with Adenine (A) and Uridine (U) led to reduced apoptosis in the most sensitive knockdown group **(Figure S4H).** These data suggest that loss of OGDHL in CRPC reduces nucleotide availability, through reduction of nucleotide synthesis and salvage genes, and increases replication stress.

### OGDHL regulates expression of neuroendocrine markers *in vitro* and *in vivo*

Prolonged AR blockade can induce changes in cell identity as tumor cells adopt AR-indifferent cell fates to adapt to treatment pressure^51^. This process, known as lineage plasticity, can give rise to aggressive and uniformly lethal molecular subtypes, such as neuroendocrine prostate cancer (NEPC)^18^. Because changes in cell fate rely on proper nucleotide balance and DNA damage repair^52,53^, we questioned whether loss of OGDHL may alter treatment-induced lineage phenotypes in prostate cancer cells. To understand how OGDHL influences response to prolonged AR blockade, we generated OGDHL knockout lines in 16D^CRPC^ cells, then treated them with Enzalutamide for 6 weeks. RNA-sequencing revealed that prolonged Enzalutamide treatment of control cells led to increased expression of genes associated with plasticity phenotypes including Axon Guidance, WNT Signaling, Epithelial-mesenchymal transition, and decreased expression of Androgen Response genes. In contrast, OGDHL knockout impaired the AR blockade-induced upregulation of plasticity-associated gene sets (**Figure 5A-5C, S5A**). Canonical indicators of lineage plasticity and antiandrogen-resistance, including upregulation of *ENO2* (encoding NSE) and *SOX9*^54^, were abrogated by loss of OGDHL (**Figure S5B, S5C**). OGDHL knockout *in vitro* led to a strong reduction in expression of *HES6* (**Figure 5D**), an evolutionarily conserved driver of neuronal development^55–57^, NE differentiation and lineage plasticity in prostate cancer^58–62^. OGDHL knockout also reduced expression of HES6 target genes^63^ (**Figure S5D**). We questioned whether this reduction in NE markers was a consequence of persistent OGDHL loss or could be induced by acute knockdown of OGDHL. We found that knockdown of OGDHL in Enzalutamide-maintained 16D^CRPC^ cells *in vitro* reduced protein expression of HES6 (**Figure S5E)** and transcription of HES6 targets (**Figure S5F),** suggesting that reduced NE marker expression is a consistent feature of OGDHL loss.

**Figure 5:**
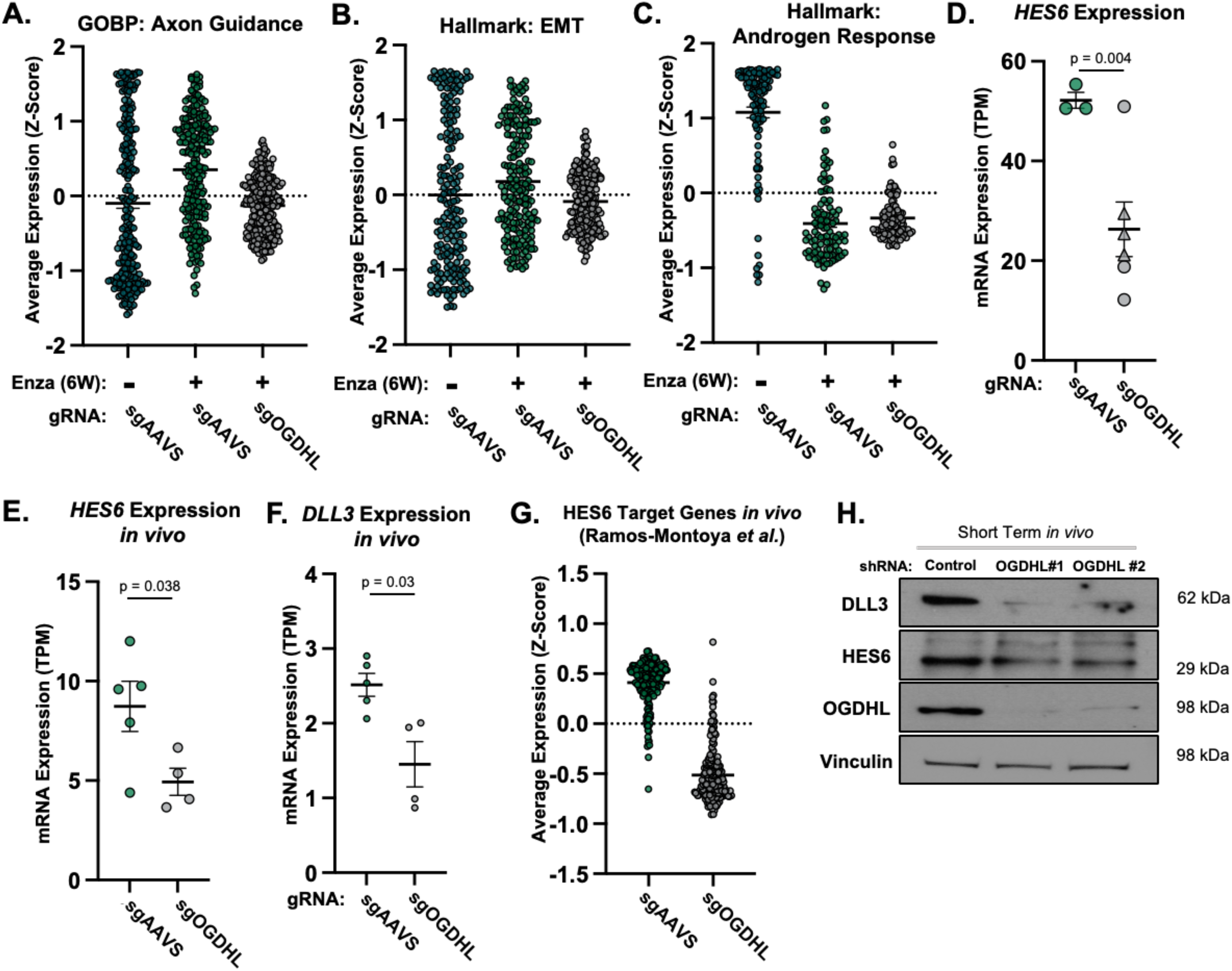
OGDHL mediates lineage plasticity phenotypes and NE marker expression in prostate cancer. **(A-C)** Average expression of genes in Gene Ontology Biological Pathway Gene Set: Axon Guidance **(A)**, Hallmark: Epithelial Mesenchymal Transition (**B)** and Hallmark: Androgen Response **(C)** gene sets in 16D^CRPC^ cells with CRISPR-Cas9 mediated genetic knockout of OGDHL allowed to adapt to enzalutamide for 6 weeks *in vitro.* Data shown is average Z scores from 3 control replicates (+/- Enza) and 6 knockout replicates (+Enza) *in vitro.* **(D)** mRNA expression of NEPC gene *HES6* in 16D^CRPC^ cells with genetic knockout of OGDHL cultured *in vitro.* **(E and F)** mRNA expression of NEPC genes *HES6* **(E)** and *DLL3* **(F)** in tumors formed from 16D^CRPC^ cells with genetic knockout of OGDHL (n =4) or control cells (n = 5). **(G)** Average expression of HES6 target genes in tumors formed from 16D^CRPC^ cells with genetic knockout of OGDHL (n =4) or control (n = 5). **(H)** Western blot of DLL3 and HES6 in control and OGDHL knockdown cell lines cells recovered 4 days after implantation *in vivo*. Error bars represent +/- SEM. P-value calculated by unpaired t-test with Welch’s Correction.

We next sought to understand if OGDHL loss could influence lineage plasticity phenotypes *in vivo.* Knockout tumors grown in castrated mice showed statistically significant reductions in transcription of NEPC genes *HES6* and *DLL3* (**Figure 5E, 5F)**. OGDHL knockout tumors also showed significantly reduced expression of HES6 target genes and other gene sets associated with AR blockade-induced plasticity phenotypes (**Figure 5G, S5G, S5H**). HES6 protein expression was also reduced following OGDHL knockdown *in vivo* (**Figure 5H, S5I, S5J)** indicating that OGDHL loss modulates expression of NEPC markers *in vitro* and *in vivo*. In summary, these data suggest that OGDHL influences AR-blockade induced lineage plasticity phenotypes.

### Mitochondrial localization is not required for OGDHL-mediated regulation of nucleotide synthesis and lineage phenotypes

Metabolic enzymes involved in various processes, including the TCA cycle, have been shown to localize to the nucleus to modulate chromatin and regulate gene expression^9–12,64–66^. As we found that OGDHL loss impacted AR blockade-induced lineage phenotypes, we attempted to revert these transitions through restoration of OGDHL. In OGDHL knockout cells, we reintroduced OGDHL with a silent mutation in the CRISPR targeted sequence. To assess the effect of mitochondrial and non-mitochondrial function, we generated both a full-length version (FL) and one lacking the mitochondrial targeting sequence (ΔMTS) (**Figure 6A**). We validated protein expression (**Figure 6A**) and the localization of each variant (**Figure 6B**). While the FL variant displayed strong colocalization with the mitochondrial marker TUFM, the ΔMTS variant displayed a diffuse, non-mitochondrial signal (**Figure 6B, S6A**). We next evaluated nuclear and mitochondrial OGDHL signal in these rescue backgrounds and the localization of endogenous OGDHL in our Enzalutamide-maintained 16D^CRPC^ cells (**Figure 6C**) via immunofluorescence. Endogenous OGDHL localization exhibited an average 40 +/- 17.5% overlap with mitochondrial signal and an average 51+/- 24.7% overlap with nuclear (DAPI) signal (**Figure 6D, 6E**). Importantly, we validated a significant reduction in mitochondrial signal, and significant increase in nuclear signal overlapping with the ΔMTS mutant compared to FL (**Figure 6D, 6E**).

**Figure 6:**
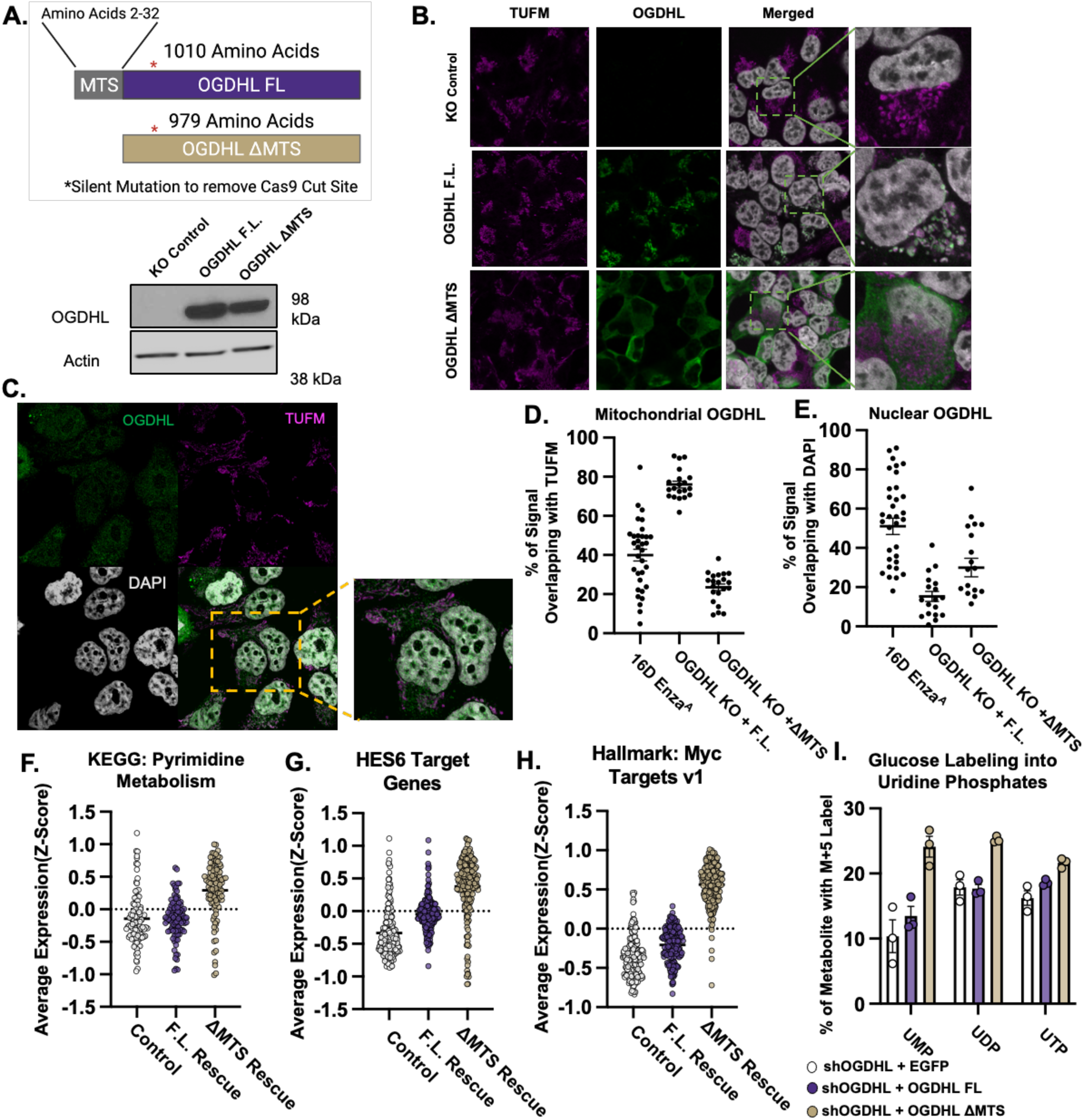
Mitochondrial localization of OGDHL is not required to regulate lineage and nucleotide phenotypes. **(A)** Schematic of CRISPR-resistant Full Length (FL) and mitochondrial targeting sequence deletion (ΔMTS) OGDHL variants and validation of protein expression. **(B)** Immunofluorescence validation of OGDHL subcellular localization in 16D^CRPC^ cells with CRISPR-Cas9 mediated genetic knockout of OGDHL and expression of knockout-resistant FL and ΔMTS variants**. (C)** Representative image of subcellular localization of endogenous OGDHL in enza-maintained 16D^CRPC^ cells. **(D and E)** Quantification of mitochondrial **(D)** and nuclear **(E)** localization of OGDHL. Data represented as % of OGDHL signal overlapping with the mitochondrial marker TUFM **(D)** or the nuclear stain DAPI **(E)**. Each data point represents a single cell. **(F-H)** Average expression of genes in the KEGG: Pyrimidine Metabolism **(F)** and HES6 Target Gene Set **(G)** and Hallmark: Myc Targets v1 **(H)** gene sets in 16D^CRPC^ cells with CRISPR-Cas9 mediated genetic knockout of OGDHL or reintroduction of knockout-resistant FL and ΔMTS variants. Data show average Z scores or TPM values from 3 technical replicates each. **(I)** M + 5 labeling of Uridine phosphates derived from U-^13^C glucose in enza-maintained 16D^CRPC^ cells with genetic knockdown of OGDHL and expression of knockdown-resistant FL and ΔMTS variants. Data represent 3 technical replicates each. Error bars represent +/- SEM. P-value calculated by unpaired t-test with Welch’s Correction.

Bulk RNA-sequencing demonstrated that restoration of OGDHL increased transcriptional expression of genes in pyrimidine metabolism, HES6 target genes, and cell cycle associated gene sets (**Figure 6F-H, S6B, & S6C**). This increase was more pronounced in cells lacking the mitochondrial targeting sequence, indicating the non-mitochondrial function of OGDHL may be more important for its impact on gene expression. At the protein level, restoration of FL and/or ΔMTS OGDHL was sufficient to abrogate OGDHL knockdown induced phenotypes including reductions in MYC and the nucleotide synthesis enzyme TK1, as well as ƔH2AX accumulation (**Figure S6D**). To assess the impact on cell metabolism, we conducted heavy-isotope glucose tracing. Consistent with our earlier findings that OGDHL does not play a meaningful role in regulating the TCA cycle, OGDHL reintroduction had limited influence on glucose incorporation into TCA cycle metabolites (**Figure S6E**). Restoration of OGDHL with either FL or ΔMTS variants increased glucose incorporation into newly synthesized pyrimidines including Uridine Phosphates (**Figure 6I**) and Cytidine Phosphates (**S6F**). Taken together, these data indicate that mitochondrial localization is not required for OGDHL-mediated regulation of lineage identity and nucleotide synthesis in prostate cancer.

### OGDHL expression is associated with neuroendocrine differentiation in prostate cancer

As we found that OGDHL loss influenced treatment-associated lineage phenotypes, we next set out to identify the expression pattern of OGDHL in advanced prostate cancer. We first interrogated TCA cycle enzyme expression in a series of murine models of prostate cancer progression driven by loss of tumor suppressors *Pten*, *Rb1* and *Tp53*^67^. Notably, the combined loss of *Pten* and *Rb1* (double knockout, DKO) or all three genes together (triple knockout, TKO) promotes a loss of luminal features and a gain of markers associated with NEPC. Among all TCA cycle genes evaluated, OGDHL was the most prominently associated with disease progression and plasticity (**Figure 7A**). OGDHL expression was greater than 10-fold higher in the NEPC-like TKO (Rb1 null;Tp53-null;PTen-null) tumors compared to the more luminal-like Pten-knockout tumors (**Figure 7B**). We next questioned whether this correlation with prostate cancer plasticity was also observed in clinical human prostate cancer and in patient-derived xenograft tumors. In RNA sequencing data collected from clinical CRPC patients^68^, we found that OGDHL expression was nearly 15-fold higher in patients with NEPC compared to CRPC-adenocarcinoma (**Figure 7C).** In the Stand Up 2 Cancer metastatic CRPC dataset^69,70^, OGDHL expression was highest in tumors with NE features, regardless of AR expression/activity, while OGDHL expression was lowest in tumors lacking NE features (**Figure 7D**). Similar results were observed in the University of Washington rapid autopsy metastatic cohort^71^ (**Figure 7E**) and in the LuCAP PDX series^72^ (**Figure 7F**) where OGDHL expression was highest in the NE+ AR-subset of tumors. Sharifi *et al.* recently defined mCRPC phenotypes based on RNA sequencing of circulating tumor cells, demonstrating that NE and LumB subtypes are associated with the shortest overall survival^73^. We found that OGDHL mRNA was detectable in CTCs and its expression was highest in the NE+ and LumB subtypes (**Figure 7G**). To ensure OGDHL was not suppressed by AR, we evaluated OGDHL expression in prostate cancer patients treated with Androgen Deprivation Therapy (ADT)^74–76^. We found that OGDHL expression was reduced in patients following ADT treatment in both paired and unpaired data sets (**Figure S7A-S7C).** Collectively, these data suggest that OGDHL is not specifically associated with AR repression but rather with plasticity and NE differentiation in CRPC.

**Figure 7:**
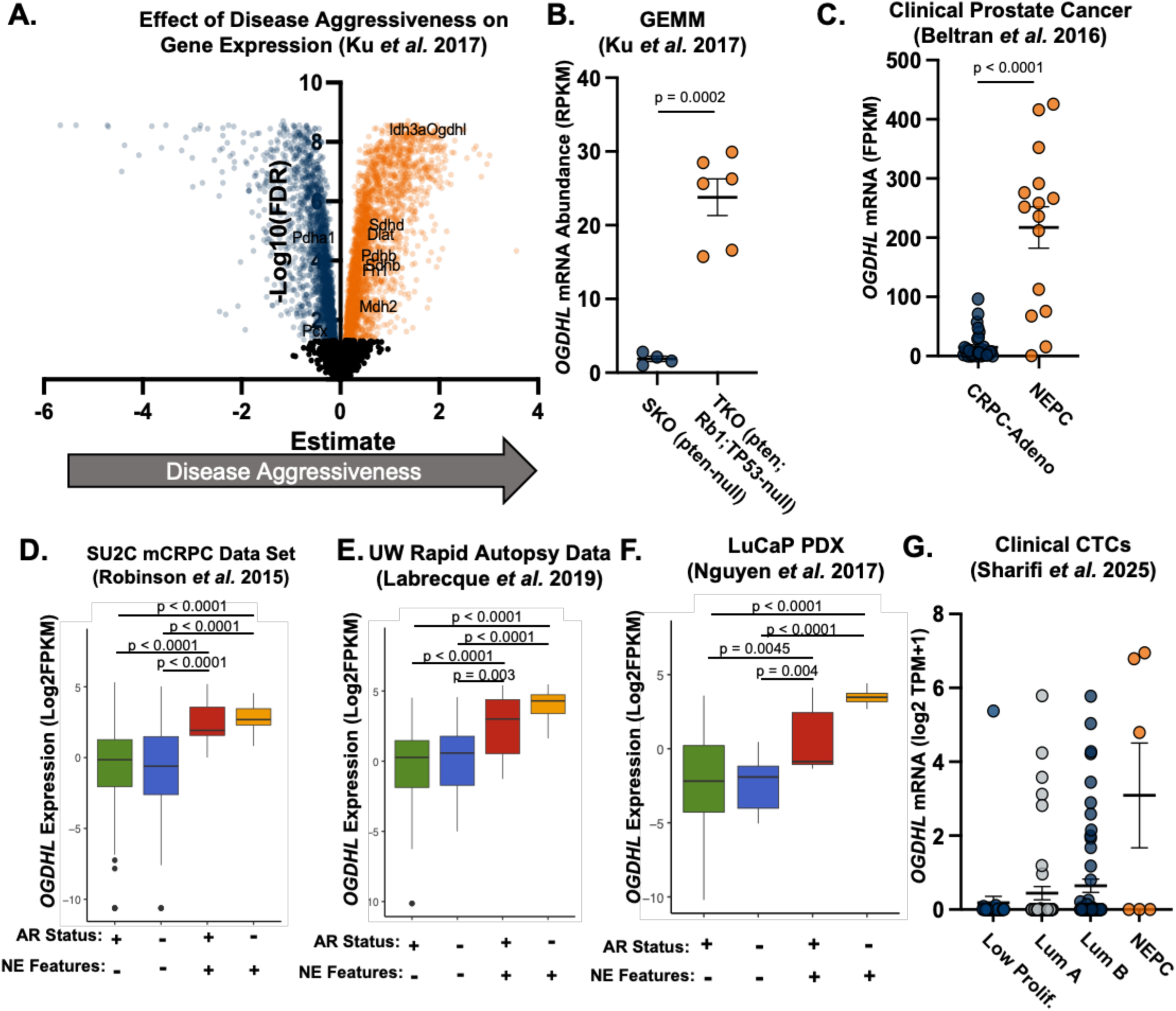
OGDHL expression is elevated in Neuroendocrine Prostate Cancer. **(A)** Volcano plot of differentially expressed genes correlated with disease progression in the Ku *et al.* GEMM data set. Highlighted genes correspond to an FDR < 0.05. Names of the top 10 differentially regulated TCA cycle genes are displayed. (**B and C)** *OGDHL* mRNA expression in a GEMM model **(B)** of prostate adenocarcinoma (pten-null (SKO)) (n=4) and NEPC (pten;Rb1;TP53-null (TKO)) (n = 6) or in patients **(C)** with Adenocarcinoma (CRPC-Adeno) or NEPC (From Beltran *et al.*). **(D-F)** *OGDHL* mRNA in clinical metastatic CRPC specimens^70^ **(D),** surgically resected tissue from patients who died from metastatic CRPC^71^ **(E)**, and in Patient Derived Xenograft models of CRPC^72^ based on AR expression and NE features^69,70^. **(G)** *OGDHL* mRNA expression in Circulating Tumor Cells (CTCs) from metastatic CRPC based on profiled subtype from the Sharifi *et al.* dataset^73.^ Error bars represent +/- SEM. P-value calculated by unpaired t-test with Welch’s Correction.

## Discussion

In this study, we demonstrate that the putative TCA cycle enzyme OGDHL is elevated at the mRNA and protein levels in response to prolonged AR blockade in CRPC cells. Despite its reported function as a tumor-suppressor gene in other cancers^30–35^, we find that OGDHL promotes antiandrogen resistant prostate cancer cell proliferation *in vitro* and *in vivo*. While some genes play a consistent role either in suppressing or driving tumorigenesis, many cancer-associated genes have context-dependent functions where they provide protective benefits in some cancer subtypes, while promoting disease progression in others. Among these are epigenetic remodelers such as EZH2 and KDM5A^77–79^, signaling factors such as TGFβ^80^, and metabolic genes such as PDHA1 and AMPK^81–83^. Our findings suggest that OGDHL represents a context-dependent regulator of cancer progression, promoting cell survival and adaptation to therapeutic pressures in prostate cancer.

Although the TCA cycle generates intermediate metabolites and biosynthetic precursors to enable metabolic and cellular functions and sustain cell growth, there is a growing appreciation for non-canonical roles of TCA cycle enzymes. Many metabolic enzymes, including the OGDH complex, have been reported to display nuclear localization to rapidly replenish metabolite pools for epigenetic reprogramming, despite their predominant expression in other subcellular compartments^9–12,64–66,84^. Metabolites, including those generated through TCA cycle enzymes, are critical substrates for epigenetic regulatory proteins, enabling the deposition and removal of methyl, acetyl, and succinyl marks on histone tails^2,8,85^. This enables cells to rapidly adapt to changes in nutrient availability, respond to extracellular signaling, and promote cell fate decisions. Because of this interconnectivity, there is a reciprocal relationship between cellular metabolism and lineage identity. OGDH complex activity modulates levels of two metabolites that are essential for faithful regulation of differentiation programs. a-KG is a critical intermediate for epigenetic regulation, as it is an obligate substrate for a class of demethylase enzymes called a-KG dependent dioxygenases (KGDDs)^86^. Alternatively, the product of OGDH complex activity, succinyl-CoA, can be used to succinylate lysine residues on histone tails, resulting in changes in chromatin accessibility and gene expression^87,88^. Here, we demonstrate that OGDHL modulates mRNA and protein expression of NEPC markers including DLL3 and HES6, suggesting that OGDHL may promote disease progression in CRPC at least in part through gene regulation. In support of that idea, modulation of OGDHL expression does not markedly alter glucose or glutamine incorporation into TCA cycle intermediates. Instead, OGDHL loss potently depletes nucleotide pools and glucose incorporation into nucleotides, while reducing expression of enzymes that fuel nucleotide synthesis and salvage.

Proper maintenance of nucleotide pools is essential to enable proliferation, gene expression, and DNA damage repair^89^. Accumulation of nucleotides is especially important for the proliferation of tumor cells due to their rapid proliferative demands, thus representing a specific vulnerability. Because of this, nucleotide metabolism is a major target for many clinically deployed cancer therapeutics^90^. However, rewiring of nucleotide metabolism in cancer can facilitate pharmacological resistance and immune cell evasion^91–93^. We find that OGDHL loss in antiandrogen-resistant prostate cancer cells depletes nucleotide pools and induces accumulation of ƔH2AX, a prominent marker of replication stress. These findings are consistent with a previous report that utilized bioinformatic approaches to implicate OGDHL as a potential DNA damage-resistance gene in prostate cancer^48^. Interestingly, OGDHL overexpression has been reported to modulate nucleotide pools and induce DNA damage in hepatocellular carcinoma, thereby inhibiting disease progression^35^. This highlights the unique function of OGDHL in prostate cancer progression. In addition to accumulation of ƔH2AX, we find that OGDHL loss reduces transcription of cell cycle associated genes and increases transcription of apoptotic and cellular stress response genes. We postulate that OGDHL is elevated in response to prolonged AR blockade in order to promote nucleotide homeostasis, which enables sustained proliferation and treatment-induced lineage plasticity. As OGDHL is not highly expressed in benign prostate tissue or localized disease, but is consistently elevated in NEPC, OGDHL may represent a novel therapeutic target for this currently uniformly lethal prostate cancer subtype.

While our findings highlight the importance of OGDHL in cell survival and lineage plasticity in prostate cancer, many questions remain. Although OGDHL modulates nucleotide metabolism, tumor growth, DNA damage response, and plasticity in CRPC, the mechanisms underlying these roles are still unclear. It is possible OGDHL plays a direct role in epigenetic regulation through enzymatic function in the nucleus, regulating a-KG and succinyl-CoA levels to modulate DNA methylation, histone methylation, and histone succinylation. As the OGDH complex does not directly modulate intermediates of *de novo* nucleotide synthesis, more direct mechanistic studies to uncover how OGDHL regulates nucleotide metabolism will be essential. OGDHL is a poorly characterized homolog of a much more broadly expressed and investigated metabolic gene, therefore more careful exploration of the nuances differentiating these two, as well as identifying targetable motifs on OGDHL, are important to clarify its potential as a therapeutic target for currently untreatable prostate cancer subtypes.

## Materials and Methods

### Materials Availability

This work did not generate new unique reagents.

### Animal Work

All animal work was performed using IACUC approved protocols under the supervision of veterinarians from the Division of Laboratory Animal Medicine at UCLA.

This method refers to figures 1K, 1L, S1J, 4G, 4H, S4E, 5H, S5I, and S5J. 750K Enzalutamide-maintained 16D^CRPC^ cells with short-hairpin (shRNA) mediated knockdown of OGDHL or a control vector were implanted subcutaneously with 20 μL of growth factor-reduced Matrigel (Corning: CB-40230C) to form tumors. After 4 days, material was collected from 4 control and 4 knockdown injections and used to generate protein lysate (Short Term *in vivo*). The remaining injections were allowed to grow for 4 weeks, after which material was collected, weighed, and used to generate protein lysate.

This method refers to figures 2F, S2F, 4A, S4A, 5E-5G, S5G, and S5H. Serial dilutions of 1M, 100K, 10K, and 1K 16D^CRPC^ cells with CRISPR-Cas9 mediated genetic knockout of OGDHL were implanted subcutaneously with 20 μL of growth factor-reduced Matrigel (Corning: CB-40230C). Once palpable tumors were measured in at least one mouse per condition (1M Cells: 6 Weeks; 100K Cells: 10 Weeks; 10K Cells: 12 Weeks, 1K cells: 16 Weeks), mice were sacrificed, tumors were resected, sectioned, and prepared as lysate or flash frozen for bulk RNA-sequencing.

This method refers to figure 1A, 1B, 1E, 1F, S1A, and S1B 550K MDa 180-30 PDX cells were implanted subcutaneously with 30 uL of growth factor-reduced Matrigel (Corning: CB-40230C). Tumors were allowed to establish for 19 days, then mice were treated with Enzalutamide or DMSO control via oral gavage daily for 14 days. *In vivo* tumor metabolism was then measured as describec below. After measurement of mitochondrial membrane potential and glucose consumption, mice were sacrificed, tumors were resected, sectioned, and prepared as lysate.

### In Vivo Assessment of Tumor Cell Metabolism by ^18^F-FBnTP and ^18^F-FDG PET/CT

Animals without fasting were warmed on a heating pad for 30 min, anesthetized with 2% isoflurane in oxygen, and injected via the tail vein with ∼90 µCi of clinical-grade ^18^F-fluorobenzyl-triphenylphosphonium (^18^F-FBnTP) or ^18^F-fluorodeoxyglucose (^18^F-FDG) in saline. Animals underwent a 1-hour conscious ^18^F-FBnTP or ^18^F-FDG biodistribution period on a heating pad prior to imaging. Exact radiotracer isotope dose, draw time, time of injection, and imaging start time were annotated to aid in signal normalization. Positron emission tomography (PET) and computed tomography (CT) scans were conducted on a G8 combined PET/CT instrument (Sofie Biosciences, Inc.) with a 600-s PET acquisition and maximum-likelihood expectation maximization reconstruction, and with a 50-s CT acquisition and Feldkamp reconstruction. PET data were converted to percent-injected dose per gram (%ID/g) and PET/CT images were co-registered. Mean values of PET signal intensity from tumor region-of-interest (ROI) normalized to heart signal uptake were analyzed using AMIDE software v1.0.4.

### Cell Lines, lentiviral transduction and cloning

Cell Lines were routinely tested for mycoplasma and authentication by short tandem repeat analysis (Laragen). Tissue culture plates were coated with 0.01% (v/v) Poly-L-Lysine Solution (Sigma-Aldrich: P4832) diluted 1:20 in ddH_2_O, then washed with sterile DPBS without Calcium and Magnesium (Gibco: 14-190-20). 16D^CRPC^ and LNCaP cells were cultured in RPMI 1640 (Gibco: 22400) supplemented with 10% (v/v) FBS (Sigma-Aldrich: F0926), and 100 units/mL penicillin-streptomycin (Gibco: 15-140-122). Media changes were conducted every 48 hours and cells were passaged to fresh tissue culture plates once they reached 90% confluency. PC3 cells were cultured in F-12K media (ATCC 30-2004) supplemented with 10% (v/v) FBS (Sigma-Aldrich: F0926), and 100 units/mL penicillin-streptomycin (Gibco: 15-140-122). LuCaP 35CR cells were cultured in Advanced DMEM F-12 supplemented with 1% Glutamax (Gibco 35050061), 10% (v/v) FBS (Sigma-Aldrich: F0926), and 100 units/mL penicillin-streptomycin (Gibco: 15-140-122). Enzalutamide treatment was performed by adding 10μM Enzalutamide (Selleck Chemicals: S1250) at each media change.

Lentivirus with short-hairpin RNA (shRNA) against OGDHL (shOGDHL) or a non-targeting control (shScrambled), Cas9 Guide RNA (gRNAs) targeting OGDHL or an AAVS control, and OGDHL overexpression vector were all obtained from Vectorbuilder (Chicago, IL, USA).

For Lentiviral transductions, cells were seeded at 30% confluence. After 24 Hours, base media containing lentivirus and 8mg/mL Polybrene (ThermoFisher Scientific: NC0663391). Successful transduction was validated by fluorescence microscopy (fluorescent vectors) or puromycin dihydrochloride selection (1.25 μg/mL) (Gibco: A1113803), 72 H after infection.

### Tumoroid Culture

This method refers to figure 1I and 1J

To generate tumoroids, cells were resuspended in a mixture of growth factor-reduced Matrigel (Corning: CB-40230C) and culture media at a ratio of 7:1 to a density of 500 cells per 80 μL. This mixture was then added to poly-hema coated 24-well culture plates (Corning: 09-761-146) as previously described^94^. Matrigel was allowed to solidify at 37°C for 60 minutes, then 350 μL of RPMI 1640 (Gibco: 22400) supplemented with 10% (v/v) FBS (Sigma-Aldrich: F0926), and 100 units/mL penicillin-streptomycin (Gibco: 15-140-122) was added to the center of the solidified ring. Media was changed every other day. After 2 weeks, at least 50 tumoroids per condition were imaged at 20X magnification and diameter was measured.

This method refers to Figure S1E

MDA-PCa 180-30^42^ PDX tumors were maintained by serial implantation of 20 – 80 mg of minced tumor tissue. Tumors were collected and cryopreserved. Upon thawing, tumor sections were mechanically dissociated and embedded in Matrigel rings to establish *ex vivo* tumoroids. Tumoroids were maintained in Human Organoid Media^95^. To passage tumoroids, media was removed and matrigel containing tumoroids was dissociated by resuspension in Advanced DMEM-F12 (Gibco: 12-634-010) containing 1 mg/mL Dispase II (Gibco: 17-105-041) and 10μM p160 ROCK-inhibitor Y-27632 2HCl (Selleck Chemicals: S1049). This mixture was incubated at 37°C for 1 hour with constant rocking. Following dissociation, cells were pelleted by centrifugation at 800xG for 5 minutes, then resuspended in prewarmed 0.05% Trypsin-EDTA, Phenol Red (Gibco: 25-300-120) and pipetted to homogenize. After 1 minute, media was added to quench the trypsinization, and cells were plated into Matrigel rings as above. To treat with enzalutamide, Human organoid media containing 10 μM Enzalutamide was added for 1 week, with fresh media supplied every other day.

### *In vitro* metabolic profiling and U-^13^C isotope tracing

This method refers to figures 3C-3F.

Enzalutamide-Maintained 16D cells harboring genetic knockdown (shRNA) OGDHL or a control vector were seeded in a 6-well plate (Corning: 07-200-83) at 325K cells/well. After 48 hours, cells were washed with PBS, then provided with glucose-free RPMI 1640 (Gibco: 11-879-020) supplemented with 11 mM U-^13^C glucose (Cambridge Isotope Labs: CLM-1396), 10% (v/v) FBS (Sigma-Aldrich: F0926), and 100 units/mL penicillin-streptomycin (Gibco: 15-140-122).

24 hours after the addition of heavy-isotope labeled media was added, cells were harvested and extracted using previously described methods^21^. In short, cells were placed on ice and washed with ice-cold 150 mM ammonium acetate, pH 7.3, then immediately treated with 500 μL of 80% MeOH (v/v) containing 10nM trifluoromethanesulfonate (internal standard). Cells were then scraped and transferred to a 1.7 mL microcentrifuge tube, vortexed 3 times on ice then centrifuged at 17,000xg for 5 minutes. The supernatant was transferred to ABC glass vials (DWK Life Sciences Wheaton 03-410-151) and dried using the EZ-2Elite Evaporator (Genevac). Samples were stored at -80°C until analysis by LC-MS.

Dried metabolites were reconstituted in 100 µL of a 50% acetonitrile (ACN) 50% dH20 solution. Samples were vortexed and spun down for 10 min at 17,000g. 70 µL of the supernatant was then transferred to HPLC glass vials. 10 µL of these metabolite solutions were injected per analysis. Samples were run on a Vanquish (Thermo Scientific) UHPLC system with mobile phase A (20mM ammonium carbonate, pH 9.7) and mobile phase B (100% ACN) at a flow rate of 150 µL/min on a SeQuant ZIC-pHILIC Polymeric column (2.1 × 150 mm 5 μm, EMD Millipore) at 35°C. Separation was achieved with a linear gradient from 20% A to 80% A in 20 min followed by a linear gradient from 80% A to 20% A from 20 min to 20.5 min. 20% A was then held from 20.5 min to 28 min. The UHPLC was coupled to a Q-Exactive (Thermo Scientific) mass analyzer running in polarity switching mode with spray-voltage=3.2kV, sheath-gas=40, aux-gas=15, sweep-gas=1, aux-gas-temp=350°C, and capillary-temp=275°C. For both polarities mass scan settings were kept at full-scan-range = (70-1000), ms1-resolution=70,000, max-injection-time=250ms, and AGC-target=1E6. MS2 data was also collected from the top three most abundant singly-charged ions in each scan with normalized-collision-energy=35. Each of the resulting “.RAW” files was then centroided and converted into two “.mzXML” files (one for positive scans and one for negative scans) using msconvert from ProteoWizard^96^. These “.mzXML” files were imported into the MZmine 2 software package^97^. Ion chromatograms were generated from MS1 spectra via the built-in Automated Data Analysis Pipeline^98^ (ADAP) chromatogram module and peaks were detected via the ADAP wavelets algorithm. Peaks were aligned across all samples via the Random sample consensus aligner module, gap-filled, and assigned identities using an exact mass MS1(+/-15ppm) and retention time RT (+/-0.5min) search of our in-house MS1-RT database. Peak boundaries and identifications were then further refined by manual curation. Peaks were quantified by area under the curve integration and exported as CSV files. If stable isotope tracing was used in the experiment, the peak areas were additionally processed via the R package AccuCor 2^99^ to correct for natural isotope abundance. Peak areas for each sample were normalized by the measured area of the internal standard trifluoromethanesulfonate (present in the extraction buffer) and by the number of cells present in the extracted well.

This method refers to Figure 3B, 3G, S3A-S3H, 6I, S6E and S6F Enzalutamide-Maintained 16D cells harboring genetic knockdown (shRNA), knockout (gRNA) of OGDHL, or a combination of knockdown vector with a shRNA resistant-OGDHL expression vector, or a matched control vector were seeded in a 6-well plate (Corning: 07-200-83) at 325K cells/well. After 48 hours, cells were washed with PBS, then provided with glucose-free RPMI 1640 (Gibco: 11-879-020) supplemented with 11 mM U-^13^C glucose (Cambridge Isotope Labs: CLM-1396), 10% (v/v) FBS (Sigma-Aldrich: F0926), and 100 units/mL penicillin-streptomycin (Gibco: 15-140-122) (for heavy-labeled glucose tracing) or glutamine-free RPMI 1640 (Gibco: 21-870-076) supplemented with 2mM U-^13^C glutamine (Cambridge Isotope Labs: CLM-1822) (for heavy-labeled glutamine tracing), 10% (v/v) FBS (Sigma-Aldrich: F0926), and 100 units/mL penicillin-streptomycin (Gibco: 15-140-122). 24 hours after the addition of heavy-isotope labeled media was added, cells were harvested and extracted using previously described methods^21^. In short, cells were placed on ice and washed with ice-cold 150 mM ammonium acetate, pH 7.3, then immediately treated with 500 μL of 80% MeOH (v/v) containing 1 nmol norvaline (internal standard). Cells were then scraped and transferred to a 1.7 mL microcentrifuge tube, vortexed 3 times on ice then centrifuged at 17,000xg for 5 minutes. The supernatant was transferred to ABC glass vials (DWK Life Sciences Wheaton 03-410-151) and dried using the EZ-2Elite Evaporator (Genevac). Samples were stored at - 80°C until analysis by LC-MS.

To calculate cell number DNA normalization was performed by resuspending the insoluble fraction in 300 uL of lysis solution (0.1M NaCl, 20mM Tris-HCL, 0.1% SDS, and 5mM EDTA in ddH_2_O). Samples were then syringed 5x using a 25G needle to homogenize. 50uL of each sample was transferred into a 96-well black-walled clear bottom plate (Corning: 07-200-588) and 100 uL of 5μg/mL Hoechst 33342 (ThermoFisher: H1399) in ddH_2_O was added. The plate was then incubated for 30 minutes at 37°C in the dark, after which DNA-based fluorescence was measured using a Tecan Infinite M1000 plate reader with 355nm excitation and 465 emission.

Dried metabolites were resuspended in 50% ACN:water and 5 ul was loaded onto a Luna 3um NH2 100A (150 × 2.0 mm) column (Phenomenex). The chromatographic separation was performed on a Vanquish Flex (Thermo Scientific) with mobile phases A (5 mM NH4AcO, pH 9.9) and B (ACN) and a flow rate of 200 μl/min. A linear gradient from 15% A to 95% A over 18 min was followed by 7 min isocratic flow at 95% A and reequilibration to 15% A. Metabolites were detected with a Thermo Scientific Q Exactive mass spectrometer run with polarity switching in full scan mode with an m/z range of 70-975 and 70.000 resolution. Maven (v 8.1.27.11) was utilized to quantify the targeted metabolites by AreaTop using accurate mass measurements (< 5 ppm) and expected retention time previously verified with standards.

Values were normalized to cell number. C13 natural abundance corrections were made using AccuCor^99^ (N15 corrections are made with AccuCor, dual-labeled corrections are made with AccuCor2). Relative amounts of metabolites were calculated by summing up the intensities of all detected isotopologues of a given metabolite. Data analysis was performed using in-house R scripts.

### Cell Viability Assay

This method refers to figures 1G, 1H, and S1I

Cell lines were detached from tissue culture plates using 0.05% Trypsin-EDTA, Phenol Red (Gibco: 25-300-120) at 37°C for 5 minutes then quenched with culture medium, then pelleted by centrifugation, followed by resuspension in base media. 10 μL of the resuspension was then mixed with Trypan Blue (Corning: 25900Cl), and cells were counted via hemocytometer.

Cells were seeded into duplicate 96-well black-walled clear bottom plate (Corning: 07-200-588) at a concentration of 10K cells/well in 100 μL of base media. Each experiment was completed with between 4 and 6 technical replicates per condition. After 16 hours, 100 μL of CellTiter-Glo® Luminescent Cell Viability Assay Reagent (Promega: G7571) was added, the plate was then incubated for 15 minutes at RT, on an orbital shaker. Luminescence was measured using a Tecan Infinite M1000 plate reader with to obtain baseline values. 48 hours later, 100 μL of CellTiter-Glo® Luminescent Cell Viability Assay Reagent (Promega: G7571) was added to the duplicate plate, then incubated for 15 minutes at RT on an orbital shaker, then luminescence was recorded. Difference in luminescence from baseline to the 48 hour timepoint was used as a proxy for changes in cell viability.

### Apoptosis Assay

This method refers to Figure S4H.

Cells were seeded in 6-well tissue culture dishes (Corning: 07-200-83) at 325K cells/well for 48H. Cell culture media and wash media were collected and pooled with quenched trypsin-containing media containing cells and apoptosis analysis was performed using an APC AnnexinV apoptosis detection kit (BioLegend, 640920) according to the provided protocol. Flow cytometry was performed to quantify the percentage of annexin V^+^ cells.

### Alkaline Comet Assay

This method refers to Figure S4F

Comet assay 3-well slides (Cell Biolabs Cat# STA-353) were pre-coated with 1.5% medium-melting-point agarose (Sigma Aldrich Cat#A6877-500g) and dried overnight. Harvested cells were resuspended in 0.5% low-melting-point agarose (Lonza Cat#50101) at 37°C at 1 to 20 ratio. Cell suspension was spread onto pre-coated slides and allowed to polymerize for 20 minutes in the dark. Then, slides were incubated in lysis buffer (10 mM Tris-HCl, pH 10, 2.5 M NaCl, 0.1 M EDTA, 1% Triton X-100) for 30 minutes. Afterwards, samples were incubated in alkaline running buffer (0.3 M NaOH, 1 mM EDTA) for 30 min before electrophoresed (300 mA, 20 V) for 15 min at 4°C. Slides were washed three times with distilled water (dH2O) and fixed with cold 70% ethanol for 15 minutes. After completely drying the slides at 37 C, stain DNA on slides using 1 uM Yoyo-1 (Fisher Cat#3601) for 30 minutes in dark. Wash and dry stained slides overnight. Images acquired using the Zeiss Axiocam 506 Mono.

### Immunoblotting

This method refers to figures 1D, 1E, S1C-S1H, S1I, S2C, S2F, 4F-4H, S4G, S4I, 5H, S5E, S5I, 6A and S6D. Cells were lysed in RIPA buffer (50 mM Tris-HCl pH 8.0, 150 nM NaCl, 1% NP-40, 0.5%Sodium Deoxycholate, 0.1% SDS) containing a phosphatase inhibitor cocktail (Halt: 78428) and a protease inhibitor cocktail (Millipore Sigma: 1169749). Sonication was performed with a sonic dismembrator (ThermoFisher Scientific: FB120). For isolation of protein from tumors, small portions were isolated using dissociation with a razor blade, then added to RIPA lysis buffer in bead mill tubes (ThermoFisher Scientific: 15-340-164). The mixture was shaken 60 seconds 2x at max intensity using a Bead Mill 4 homogenizer (ThermoFisher Scientific: 15-340-164).

Protein concentration was calculated by the Pierce™ BCA Protein Assay (ThermoFisher Scientific: PI23228) according to manufacturer specifications. Samples were run on NuPAGE 4-12% Bis-Tris Gels (ThermoFisher Scientific: NP0335, NP0322) at 200V for 40-50 minutes, then transferred to Immobilon™-P PVDF Membranes (Millipore Sigma: IPVH 00010) at 30V for 60 minutes. Total protein was visualized using the Sypro Ruby protein blot stain (ThermoFisher Scientific: S11791).

Membranes were blocked in 5% Milk (LabScientific: M0841) in DPBS without calcium or magnesium (Gibco: 21-600-044) with 0.1% Tween-20 (Fisher: BP337) for at least 1 hour at RT on an orbital shaker. Membranes were then probed with primary antibodies overnight at 4°C, followed by chromophore-conjugated Alexa Fluor 647 anti-mouse (ThermoFisher Scientific: A21235), Alexa Fluor 647 anti-rabbit (ThermoFisher Scientific: A21244), HRP-conjugated anti-mouse (ThermoFisher Scientific: 31437), or anti-rabbit (ThermoFisher Scientific: G-21234) secondary antibodies. Chromophore-conjugated antibodies were detected by fluorescence using a GE Typhoon FLA 9000 Gel Imaging Scanner or Invitrogen IBright 1500. HRP-conjugated antibodies were detected using chemiluminescent detection on film (Fisher: PI34091). Primary antibodies used were β-Actin (Invitrogen: PIMA-1140), Prostate Specific Antigen (PSA) (Cell Signaling: 5877S), OGDHL (Invitrogen MA5-62626), OGDH (Proteintech: 15212-1-AP), Phospho-Histone H2A.X(Ser 139) (ƔH2AX) (Cell Signaling: 9718S), Vinculin (Abcam: ab129002), HES6 (NovusBio: NBP3-04548), and DLL3 (Cell Signaling: 71804S).

### *In vitro* RNA Sequencing

This method refers to Figure 2A-E, S2A, S2B, S2D, S2E, 4B-4E, S4B-S4D, 5A-5D, S5A-S5D, 6F-6H, S6B and S6C.

Cells were plated into 6-well tissue culture dishes (Corning: 07-200-83) at 325K cells/well. After 48 hours, cells were lifted cell lines were detached from tissue culture plates using 0.05% Trypsin-EDTA, Phenol Red (Gibco: 25-300-120) at 37°C for 5 minutes then quenched with culture medium, then pelleted by centrifugation. Supernatant was removed and cells were washed once with sterile DPBS without Calcium and Magnesium (Gibco: 14-190-20), then the supernatant was removed, and cells were flash frozen using Liquid N_2_. Cell pellets were maintained at -80°C until submission for sequencing.

### *In vivo* RNA Sequencing

This method refers to Figure 2F, 4A, S4A, 5E-5G, S5G, and S5H

Small tumor sections were isolated from harvested tumors and flash frozen using flash frozen using Liquid N_2_. Tumor sections were maintained at -80°C until submission for sequencing.

Libraries for RNA-Seq were prepared with KAPA mRNA HyperPrep Kit. The workflow consists of mRNA enrichment and fragmentation, first strand cDNA synthesis using random priming followed by second strand synthesis converting cDNA:RNA hybrid to double-stranded cDNA (dscDNA), and incorporates dUTP into the second cDNA strand to maintain strand origin information. cDNA generation is followed by end repair to generate blunt ends, A-tailing, adaptor ligation and PCR amplification. Different adaptors were used for multiplexing samples in a half lane. Sequencing was performed on Illumina NovaSeq X Plus for PE 2x50 run. Data quality check was done on Illumina SAV. Demultiplexing was performed with Illumina Bcl2fastq v2.19.1.403 software. The alignment was performed using STAR^100^ with human reference genome GRCh38. The Ensembl Transcripts release GRCh38.107 GTF was used for gene feature annotation. For normalization of transcripts counts, TPM normalized counts were generated.

### Confocal Microscopy

This method refers to Figure 6B and 6C

Cells were plated onto sterilized glass coverslips in 24-well tissue culture dishes (Corning: 09-761-146) at 15K cells/well. Once cells reached 80% confluency, media was removed and cells were washed with sterile PBS. Cells were then fixed in 4% Paraformaldehyde in PBS for 20 minutes at RT. After fixation, cells were washed with PBS, then permeabilized using 0.5% Triton X-100 in PBS for 5 minutes at 4°C, followed by 3 PBS washes. Coverslips were then blocked with PBS containing 3% BSA and 0.1% Tween-20 for 30 minutes at RT, before adding primary antibody for 1 Hour at RT in the dark. Following primary antibody staining, coverslips were washed 3 times with PBS containing 0.1% Tween-20. Then fluorophore-conjugated secondary antibodies were added for 1 hour at RT in the dark. Following secondary staining, coverslips were washed twice with PBS containing 0.1% Tween-20. DAPI (1:2000) was added during a final wash in 0.1% Tween-20 for 5 minutes. Coverslips were mounted onto glass slides using Prolong Gold (Thermofisher: P36930) and allowed to dry overnight in the dark. Slides were sealed with clear nail polish until imaging. Imaging was completed using the Zeiss LSM 980 Airyscan 2 using a 63X oil immersion objective. Primary Antibodies used were OGDHL (Invitrogen MA5-62626) and TUFM (Atlas: AMAB909066), Secondary Antibodies used were anti-Rabbit IgG Alexa Fluor 647 and anti-Mouse IgG Alexa Fluor 594.

### Differential gene expression models

This method refers to Figure 7A

Differentially expressed gene (DEG) models were fit in R (v4.2.3)(R Core team) using the limma package (v3.54.2) lmFit function^101^. Models were constructed using the log2-transformation of reads per kilobase of transcript per million reads mapped (RPKM) plus one pseudo count as the outcome and an ordinal encoding of genotype as the predictor for each gene independently as follows:

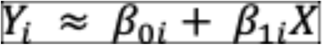

Here *Y_i_* represents a sample outcome vector for an individual gene and *β_oi_* represents the model intercept for an individual gene. *X* represents the genotype vector, which is defined as a z-scored vector of integers representing the varying genotypes in order of disease severity being experimentally modeled. *β_1i_* represents the multiplicative change (in doublings) in expression per standard deviation increase in genotypic sate for an individual gene.

Standard errors were moderated using the limma eBayes function^101^. Estimate P-values were adjusted using the Benjamini-Hochberg false discovery rate (FDR) procedure.

### RNA Alignment and Quality Control

This method refers to Figure S7A-S7C.

RNA sequencing data, provided as FASTQ files, were obtained for multiple study cohorts (Long et al., NCT01990196, Sharma et al., Wilkinson et al.)^74–76^. All datasets were processed uniformly through metapipeline-RNA (v1.0.0)^102^. Reads were aligned to the GRCh38.14 human reference genome using the STAR aligner (v2.7.11a)^101^. Gene annotations were based on GENCODE Release 45^103^. Initial quality control (QC) assessments for the raw sequencing reads were conducted using FastQC (v0.11.8), which evaluated metrics such as per-base sequence quality, sequence content, and GC content. Post-alignment QC metrics were generated by STAR, including error rates, read length distributions, splicing characteristics, and overall alignment statistics (e.g., percentage of uniquely mapped reads). MultiQC (v1.25) was employed to aggregate and summarize the QC metrics from both FastQC and STAR across all samples^104^. Samples identified as QC outliers using a significance cutoff of 0.05 based on FastQC statistics were removed using OmicsQC^105^. Subsequently, gene and transcript quantification was performed using RSEM (v1.3.3) based on the STAR alignments^106^

### Statistical Analysis of OGDHL Gene Expression in patients pre- and post-ADT Therapy

Gene expression levels for *OGDHL* (Oxoglutarate Dehydrogenase L), quantified as Transcripts Per Million (TPM), were used for differential expression analysis. Expression values were log_2_-transformed after adding a pseudocount of 1 (log_2_(TPM + 1)) prior to statistical testing and visualization. Data from the individual cohorts were aggregated for combined analyses where appropriate.

To compare *OGDHL* expression between ‘Pre’ and ‘Post’ conditions using all available samples irrespective of pairing, a two-sided Wilcoxon rank-sum test was performed on the aggregated dataset (n=231 Pre, n=109 Post samples; Fig. S7A).

For samples where paired data were available (i.e., measurements from the same individual under both ‘Pre’ and ‘Post’ conditions, n=84 pairs), a two-sided Wilcoxon signed-rank test was employed to assess significant changes in expression within individuals (Fig. S7B). In cases where multiple samples from one condition were present within an individual, their average TPM was taken and considered as a single sample.

The log_2_ Fold Change (log_2_FC) was calculated for each study cohort and the combined dataset by comparing the average log_2_(TPM + 1) expression in the ‘Post’ condition relative to the ‘Pre’ condition.

A summary dot plot was generated to visualize the differential expression results across studies (Fig. S6C). In this plot, the size and color of the dots represent the calculated log_2_FC. The statistical significance of the comparison for each study (and the combined ‘All’ dataset), stratified by unpaired and paired analysis types, is indicated by the background shading intensity. P-values obtained from the respective statistical tests (Wilcoxon rank-sum for unpaired, Wilcoxon signed-rank for paired) were adjusted for multiple comparisons across the studies shown using a Bonferroni correction. Darker shading corresponds to a lower adjusted P-value.

All statistical analyses and data visualizations were performed using R version (v4.3.3), BoutrosLab.plotting.general (v7.1.2), data.table (v1.17.0), and cowplot (v1.1.2)^107–110^.

### Statistical Analysis

#### Western Blot Quantification

Relative expression was calculated by opening image files in FIJI then inverting to negative. Mean band intensity was measured and plotted as a ratio of protein of interest/loading control. P-values were calculated using unpaired t-tests with Welch’s correction.

#### Statistical analysis of tumor formation rate

Equivalent volume of injection media was weighed as a baseline for input material. Tissue recovered after 4 weeks *in vivo* was weighed and compared to the injection media baseline. P-values for tumor formation was calculated by a 2-tailed Fisher’s Exact Test.

#### Metabolomics Analysis

Metabolite abundance and isotopologue distributions were calculated by normalizing to cell number. Metabolite abundance was measured as the sum of all isotopologues. Fold changes were calculating by averaging metabolite abundance values of control cells relative. False Discovery Rate (FDR) was calculated by applying the Benjamini-Hochberg procedure for multiple hypothesis testing to p-values corresponding to 2-tailed t-tests. Volcano plot was generated by plotting the log_2_ transformed fold change and negative log_10_ transformed FDR values. Heat maps were generated by plotting Row Z scores of metabolite abundance of knockdown cells or control.

#### RNA sequencing Analysis

Fold changes in RNA expression were calculated by normalizing measured TPM values to the average of the control group. False Discovery Rate (FDR) was calculated by applying the Benjamini-Hochberg procedure for multiple hypothesis testing to p-values corresponding to 2-tailed t-tests. Heat maps were generated by plotting fold change or log_2_ transformed fold change in TPM relative to control. Volcano plot was generated by plotting the log_2_ transformed fold change and negative log_10_ transformed FDR values. Gene Set Enrichment Analysis (GSEA) was performed using the Hallmark gene sets from the Molecular Signatures Database (MSigDB) using GSEA (v 4.3.3)^111,112^. Gene sets were gathered from the MSigDB or specified source. Row Z scores were calculated, then plotted as the average for the control or OGDHL knockdown/knockout condition.

Previously Published Data Sets

RNA-sequencing data sets were generated as previously described^67–73^. Bulk RNA-sequencing data that was re-analyzed for this manuscript were obtained as normalized transcript values. Figures 6A and 6B were generated from data published by Ku *et al.*^67^. Figure 6C was generated by NEPC patient tumor data published by Beltran *et al.*^68^. Data for Figures 6D-6E were collected from the StandUp2Cancer mCRPC dataset^69^, UW Rapid Autopsy Data Set^71^, and the LUCaP PDX Data Set^72^, stratification of groups completed by Labraque *et al.*^69^. Data for Circulating Tumor Cells obtained from Sharifi *et al.*^73^ Statistical analysis was completed by unpaired T-Test with Welch’s correction.

#### Immunofluorescence Quantification

Mitochondrial colocalization analysis was completed by opening TUFM (mitochondrial) and OGDHL channels using FIJI then measured by the Colocalization Function. OGDHL localization was calculated by measuring total OGDHL Fluorescence intensity in FIJI, then subtracting out the Mitochondrial (TUFM) or Nuclear (DAPI) signal regions using Region of Interest (ROI) Manager. Background values were removed by subtracting the Mean Fluorescence observed in OGDHL knockout cells.

## Data Accessibility

RNA-Sequencing files were uploaded to Gene Expression Omnibus: Accension IDs: GSE298123, GSE297510, and GSE298381

*Raw metabolomics data is is available at the NIH Common Fund’s National Metabolomics Data Repository (NMDR) website, the Metabolomics Workbench*^113^, https://www.metabolomicsworkbench.org *where it has been assigned Project IDs 5929 and 5931*.

## Acknowledgements

M.J.B. acknowledges the support of the Ruth L. Kirschstein National Research Service Award GM007185 and the NIDDK TL1 DK132768 Award. J.A.D. is supported by the Eugene V. Cota-Robles Fellowship. R.A. is supported by the UCLA-Caltech Medical Scientist Training Program T32GM152342 and the Jonsson Comprehensive Cancer Center Fellowship. S.L. is supported by the UCLA Jonsson Comprehensive Cancer Center. M.N.S is supported by the 2022 Point Biopharma Young Valor Investigator award and the Department of Defense PC220240. M.C.H. is supported by the NIH/NCI (R37CA286450), Grant 2021184 from the Doris Duke Charitable Foundation, the V Foundation, the Prostate Cancer Foundation Felix Feng PC-SYNERGY award and the UW/FHCC Institute for Prostate Cancer Research. D.B.S. is supported by NIH/NCI grants R01CA267721 and R01CA208642. P.C.B. is supported by the NIH grants P30CA016042, U2CCA271894, R01CA270108 and Department of Defense grants W81XWH2210247 and W81XWH2210751. A.S.G. is supported by UCLA Prostate Cancer Specialized Programs of Research Excellence (SPORE) NCI P50 CA092131, Department of Defense PCRP award HT94252310379, the Mike Slive Foundation for Prostate Cancer Research, the UC Cancer Research Coordinating Committee (C23CR5598), the 2024 Larry & Sherry Benaroya-Prostate Cancer Foundation Challenge Award, the Basser Center for BRCA, the UCLA Eli and Edythe Broad Center of Regenerative Medicine and Stem Cell Research Rose Hills Foundation Innovator Grant, the UCLA Jonsson Comprehensive Cancer Center and Eli and Edythe Broad Center of Regenerative Medicine and Stem Cell Research Ablon Scholars Program, the National Center for Advancing Translational Sciences UCLA CTSI Grant UL1TR001881, STOP CANCER, and the UCLA Institute of Urologic Oncology. The authors would like to thank Dr. Peter Nelson for LuCAP35CR cells and Dr. Nora Navone for MDA-PCa 180-30 patient derived xenograft tissue. We thank the UCLA metabolomics core and members of the Christofk lab for assistance with mass spectrometry and guidance in experimental design. We would also like to thank the UCLA Technology Center for Genomics & Bioinformatics (TCBG) for help with RNA sequencing and data analysis. *This work is supported by Metabolomics Workbench/National Metabolomics Data Repository (NMDR) (grant# U2C-DK119886), Common Fund Data Ecosystem (CFDE) (grant# 3OT2OD030544) and Metabolomics Consortium Coordinating Center (M3C) (grant# 1U2C-DK119889). Some figures were created using BioRender: Created in BioRender. Bernard, M. (2025)* https://BioRender.com/82da7ku. The funders had no role in study design, data collection and analysis, decision to publish or preparation of the manuscript. We also thank Bill Lowry, Leigh Ellis, Hilary Coller, and Brigitte Gomperts for providing critical feedback and intellectual support during the project.

## Author Contributions

M.J.B., A.R., J.A.D., R.N.D., K.Y.H., S.B., A.G., E.L., S.Z., and T.H. conducted the experiments. M.J.B., N.M.N, C.M.S., and R.A. completed data quantification and statistical analyses. M.J.B. and A.S.G. designed the experiments, wrote, and edited the manuscript. B.R.W. performed molecular cloning and advised on genetic knockout experiments. N.M. and H.R.C. provided metabolomics expertise, performed mass spectrometry on *in vitro* nutrient tracing experiments, and wrote the related methods section. S.L. and D.B.S provided expertise with PET experiments. N.M.N, R.A., H.M.R., M.C.H. and P.C.B. analyzed clinical data sets and provided bioinformatics expertise. M.N.S., J.M.L., S.G.Z., analyzed and provided data corresponding to circulating tumor cells. A.S.G. procured funding and supervised the project.

