## Supplementary Figures for "OGDHL regulates tumor growth, neuroendocrine marker expression, and nucleotide abundance in prostate cancer"

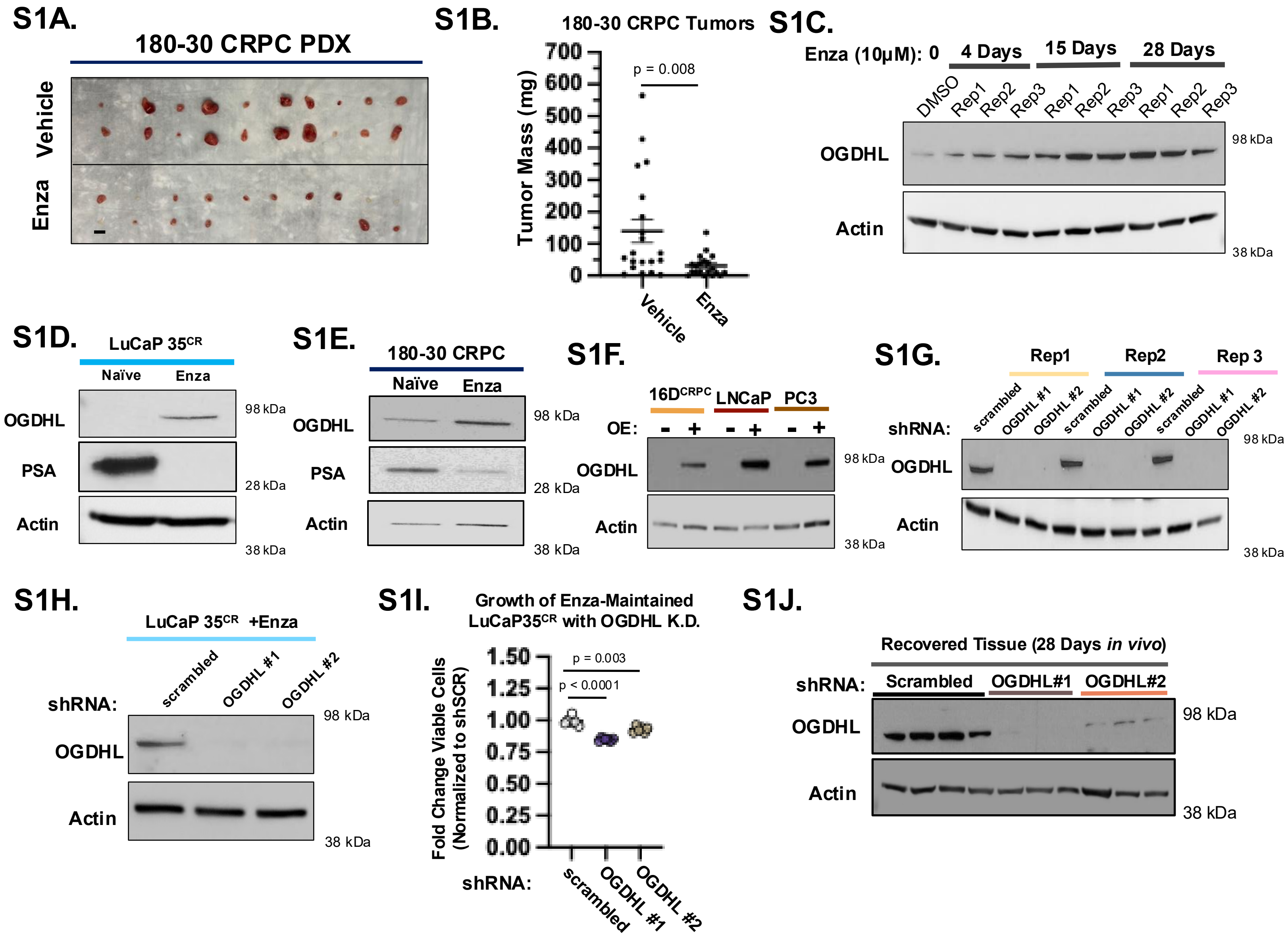


**Supplementary Data 1: Validation of Altered OGDHL Expression and reduction in growth *in vitro* and *in vivo***

**(A and B)** Images **(A)** and measured tumor weights **(B)** of MDA-PCa 180-30 CRPC Xenograft Tumors treated with vehicle or enzalutamide for 14 days by oral gavage. Scale Bar = 1.0 cm. **(C - E)** Increase in OGDHL expression with enzalutamide treatment in a time course in 3 separately maintained biological replicate $16D^{CRPC}$ cell lines **(C),** LuCaP $35^{CR}$ cells **(D),** and MDA-PCa 180-30 organoids **(E).** **(F)** Validation of successful overexpression of OGDHL in $16D^{CRPC}$, LNCaP, and PC3 prostate cancer cell lines. **(G and H)** Validation of reduced OGDHL expression in cells transduced with OGDHL knockdown vectors in 3 separately maintained biological replicate 16D lines **(G)** and Enza-maintained LuCaP $35^{CR}$ cells **(H).** **(I)** Relative change in cell viability over 48 hours of Enza-Maintained LuCaP $35^{CR}$ cells transduced with OGDHL knockdown vectors. Plotted as change in viability relative to control cells. Data from 5 technical replicates. **(J)** Western Blot validation of reduced OGDHL expression in tissues recovered 4 weeks after implantation *in vivo* from Enza-maintained $16D^{CRPC}$ cells with control (Scrambled) or OGDHL knockdown.

Error bars represent +/- SEM. P-value calculated by unpaired t-test with Welch's Correction

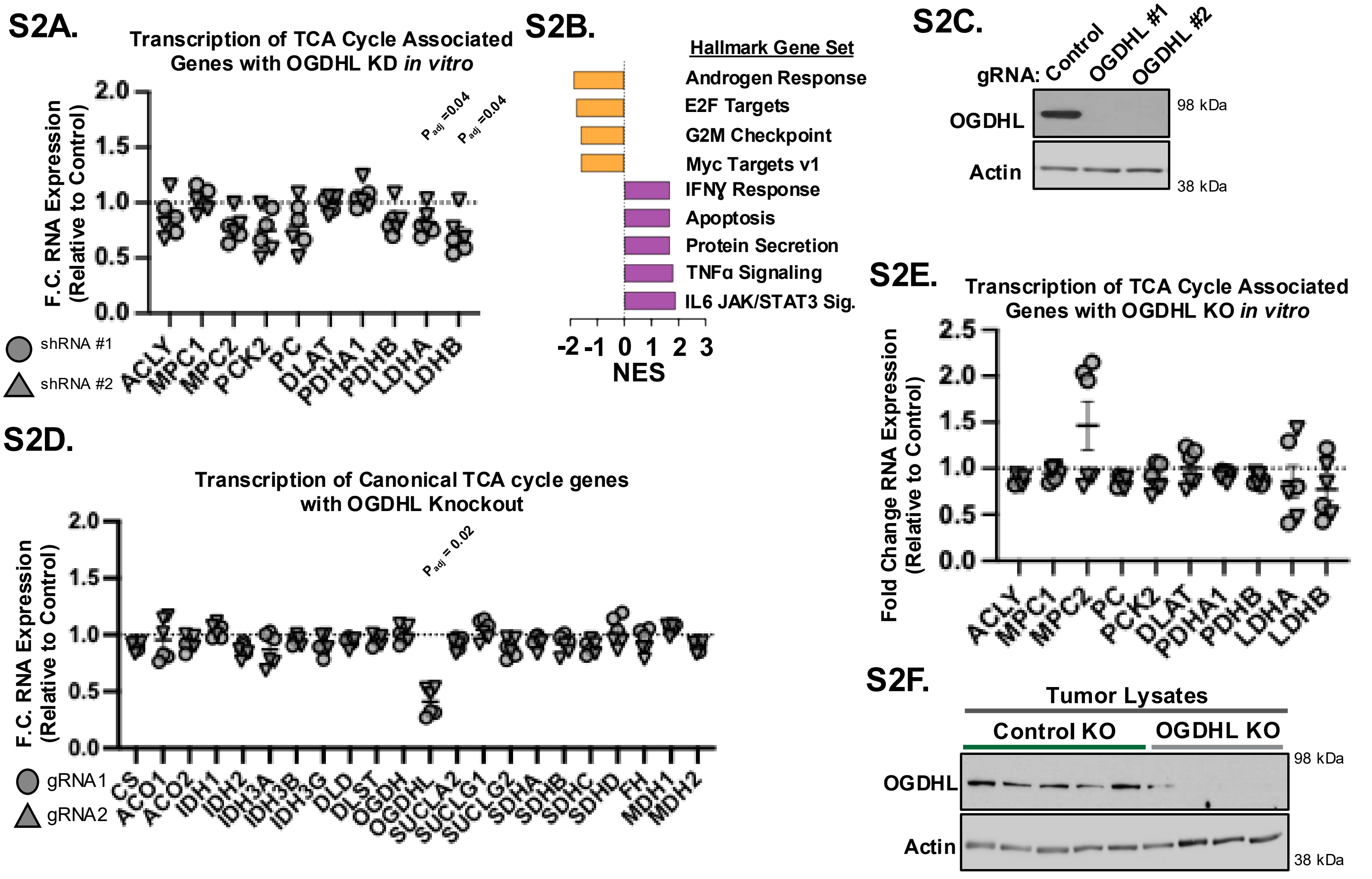


**Supplementary Data 2: Effect of OGDHL loss on expression of TCA-cycle associated genes and validation of CRISPR knockout**

**(A)** RNA expression of TCA-associated genes with OGDHL knockdown in 16D$^{CRPC}$ cells relative to control knockdown cells *in vitro.* Data represent 3 technical replicates each from 2 shRNAs. **(B)** Waterfall plot of GSEA scores for hallmark gene sets significantly (FDR < 0.05) enriched (Purple) or depleted (orange) in Enza-maintained 16D$^{CRPC}$ cells with OGDHL knockdown relative to control knockdown cells. **(C)** Western blot validation of reduced OGDHL expression in cells transduced with OGDHL CRISPR Knockout vector. (**D and E**) RNA expression of canonical TCA cycle genes **(D)** and TCA cycle-associated genes **(E)** in OGDHL knockout 16D$^{CRPC}$ cells relative to control knockout cells *in vitro.* Data represent 3 technical replicates each from 2 gRNAs. (**F**) Western Blot validation of reduced OGDHL expression in tumors generated from OGDHL knockout (KO) 16D$^{CRPC}$ cells.

Error bars represent +/- SEM. Adjusted P values were calculated by applying the Benjamini-Hochberg procedure for multiple hypothesis testing to p-values corresponding to 2-tailed t-tests. Statistically significant values shown

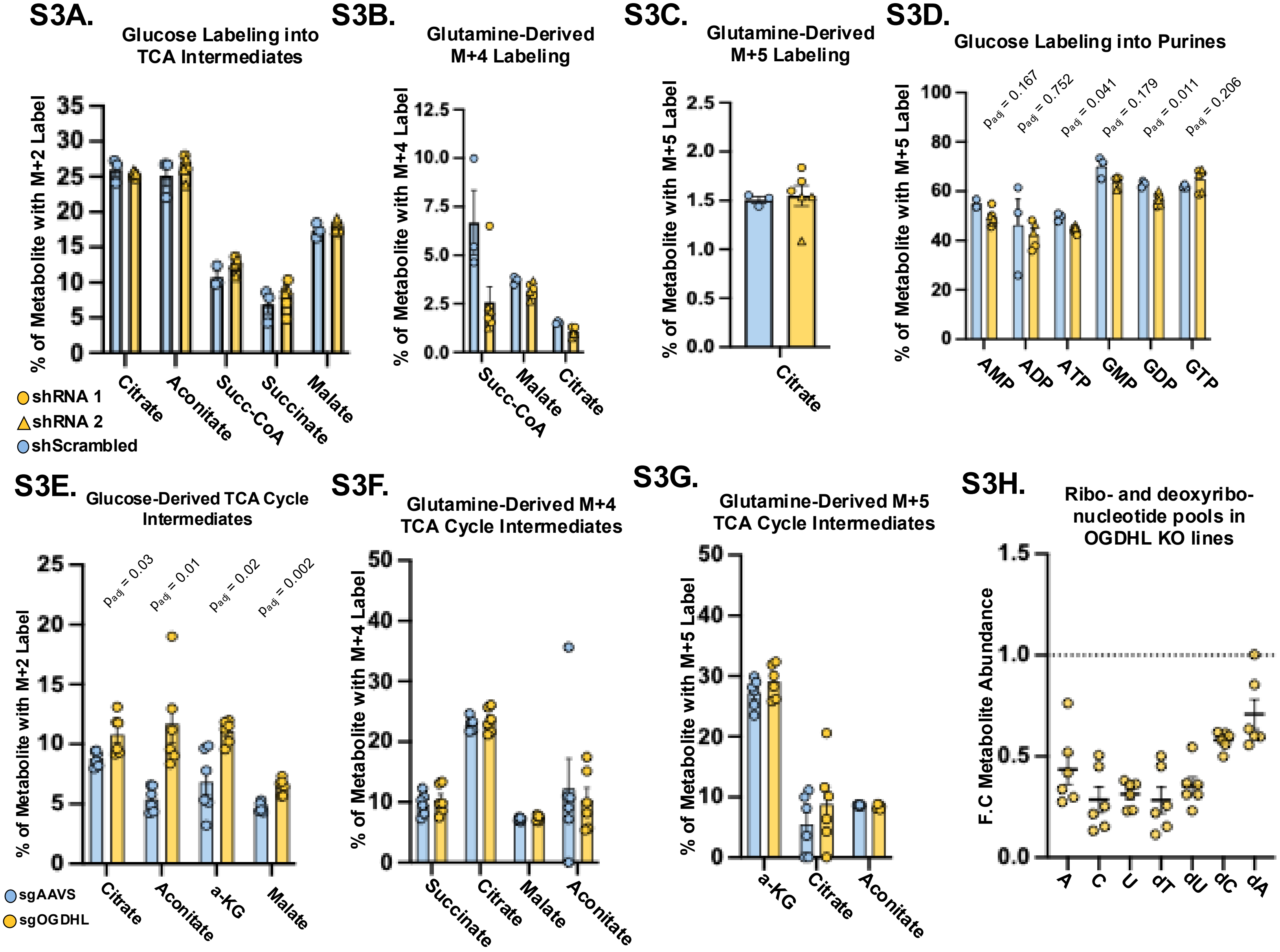


**Supplementary Data 3: OGDHL Loss in CRPC cells reduces abundance of key nucleotide synthesis precursors but does not alter glucose or glutamine incorporation into TCA cycle intermediates**

**(A-D)** M + 2 labeling of TCA cycle intermediates from U-$^{13}$C Glucose incorporation **(A)**, M + 4 labeling of TCA cycle intermediates from U-$^{13}$C glutamine incorporation through oxidative metabolism of α-Ketoglutarate **(B)** and M + 5 Labeling of TCA cycle intermediates from U-$^{13}$C glutamine incorporation through reductive carboxylation of α-Ketoglutarate **(C)** M + 5 labeling of purine phosphates derived from U-$^{13}$C glucose **(D)** in control and OGDHL knockdown Enza-maintained 16D$^{CRPC}$ cells**. (E-G)** M + 2 labeling of TCA cycle intermediates from U-$^{13}$C Glucose incorporation into TCA cycle intermediates **(E)**, M + 4 labeling of TCA cycle intermediates from U-$^{13}$C glutamine incorporation through oxidative metabolism of α-Ketoglutarate **(F),** and M + 5 Labeling of TCA cycle intermediates from U-$^{13}$C glutamine incorporation through reductive carboxylation of α-Ketoglutarate **(G)** in Enza-maintained 16D$^{CRPC}$ cells with genetic knockout of OGDHL. Data represents 3 technical replicates each from 2 gRNAs. **(H)** Abundance of ribonucleotides and deoxyribonucleotides in OGDHL knockout Enza-maintained 16D$^{CRPC}$ Cells relative to control knockout lines. Dashed line represents equivalent abundance.

Error bars represent +/- SEM. Adjusted P values were calculated by applying the Benjamini-Hochberg procedure for multiple hypothesis testing to p-values corresponding to 2-tailed t-tests.

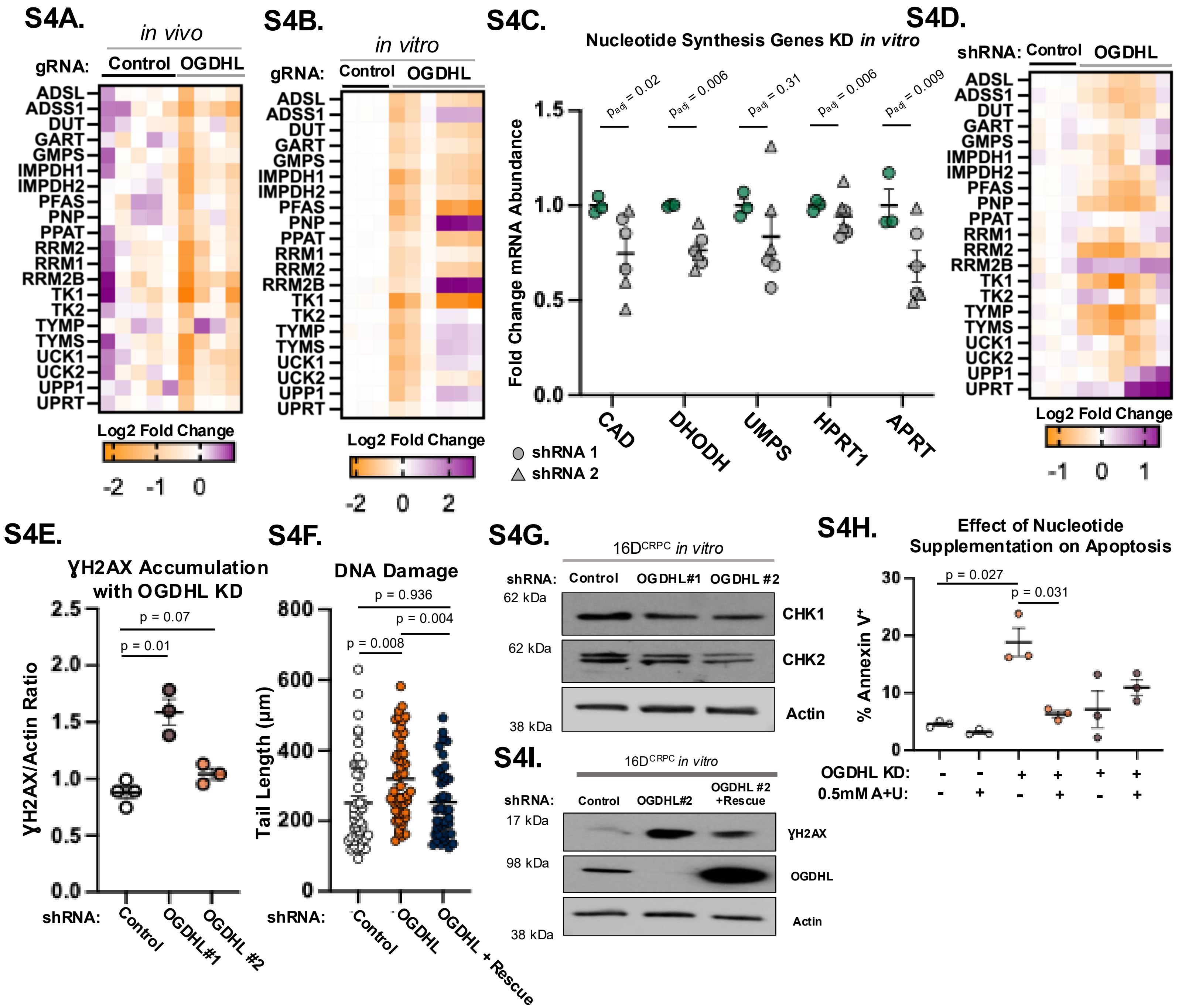


**Supplementary data 4: OGDHL Loss reduces nucleotide synthesis genes and induces hallmarks of DNA repair stress** (**A and B)** Heatmap of relative RNA abundance of nucleotide metabolism genes in tumors formed from 16D$^{CRPC}$ cells with genetic knockout of OGDHL (n =4) or a control (n = 5) **(A)** or in OGDHL knockout cells cultured *in vitro* **(B).** Data from 3 technical replicates each from 2 gRNAs. **(C and D)** Relative mRNA abundance of nucleotide synthesis genes **(C)** and nucleotide metabolism genes **(D)** in 16D$^{CRPC}$ cells with genetic knockdown of OGDHL or a control. Data from 3 technical replicates each from 2 shRNAs. **(E)** Quantification of DNA damage indicator ϒH2AX in control and OGDHL knockdown cells recovered 4 weeks after implantation *in vivo*, from Figure 4H. Data represented as the relative band intensity normalized to Actin loading control. **(F)** Quantification of DNA tail length from the fluorescence-based single-cell gel electrophoresis Comet Assay. Data represented as the tail length from individual cells: control n = 46; OGDHL Knockdown n = 51, and OGDHL Knockdown with shRNA resistant OGDHL (Rescue) n = 40. **(G)** Western blot of the DNA damage repair markers CHK1 and CHK2 in control and OGDHL knockdown cell lines *in vitro.* **(H)** Measured percentage of cells expressing the Apoptotic cell marker Annexin V, as measured by Flow Cytometry. Data from 3 technical replicates each from 2 shRNAs, in the presence or absence of 0.5mM A and U. **(I)** Western blot of the DNA damage marker ϒH2AX and OGDHL expression in control, OGDHL knockdown, and OGDHL knockdown with shRNA resistant OGDHL (Rescue) cells *in vitro.*

Error bars represent +/- SEM. P-value calculated by unpaired t-test with Welch's Correction

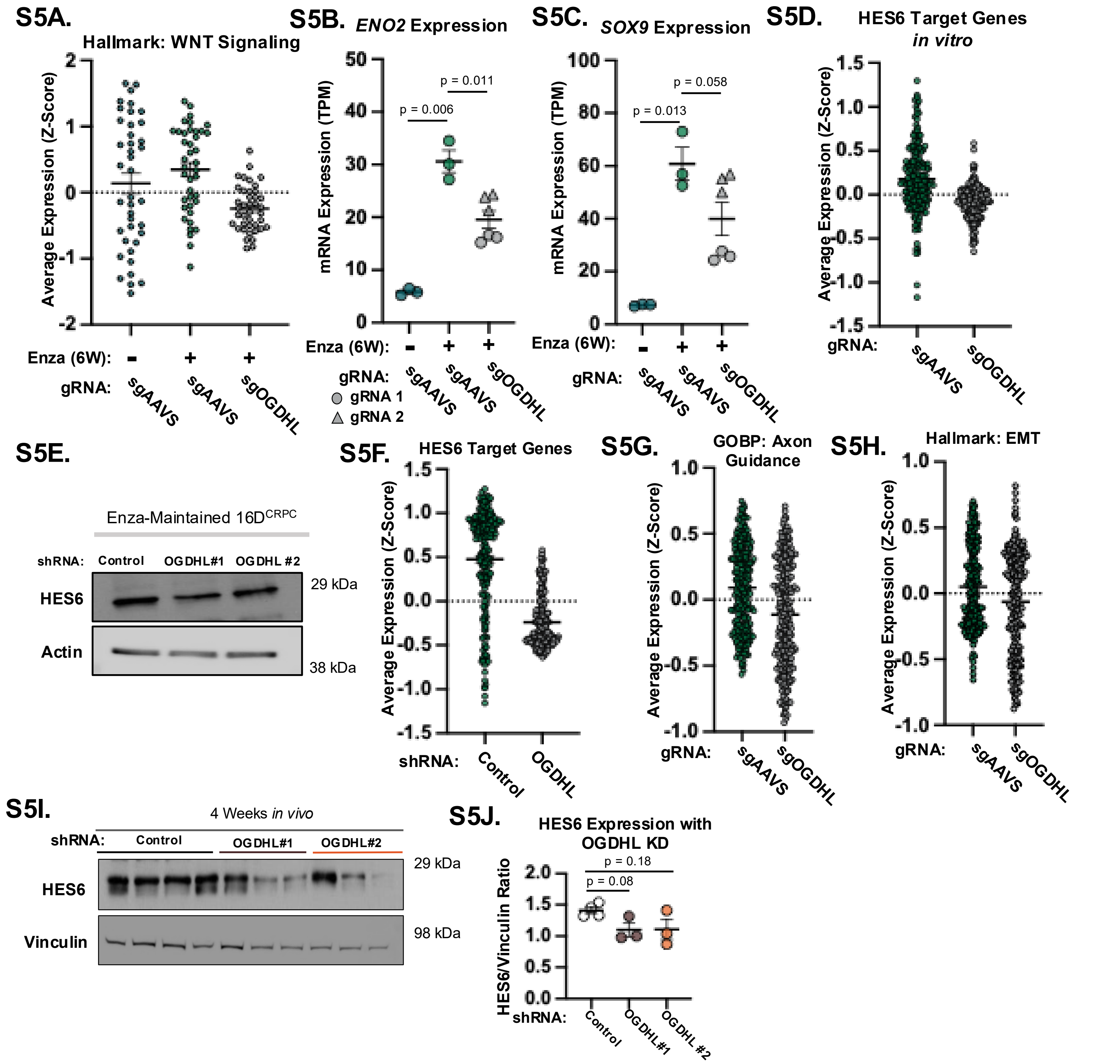


**Supplementary Data 5: OGDHL mediates lineage plasticity phenotypes and NE marker expression in prostate cancer**
**(A-C)** Average expression of genes in the Hallmark: WNT Signaling gene sets **(A)** or expression of NEPC-associated genes gene *ENO2* **(B)** and *SOX9* **(C)** in 16D$^{CRPC}$ cells with CRISPR-Cas9 mediated genetic knockout of OGDHL allowed to adapt to enzalutamide for 6 weeks *in vitro.* Data shown is average Z scores or TPM values from 3 control replicates (+/- Enza) and 6 knockout replicates (+Enza) *in vitro.* **(D)** Average expression of genes in the HES6 Target Gene Set (Curated by Ramos-Montoya *et al.* 2014) in 16D$^{CRPC}$ cells with CRISPR-Cas9 mediated genetic knockout of OGDHL. Data shown is average Z scores from 3 control replicates and 6 knockout replicates *in vitro.* **(E)** Western Blot of NEPC driver HES6 in control and OGDHL knockdown cell lines *in vitro.* **(F)** Average expression of genes in the HES6 Target Gene Set (Curated by Ramos-Montoya *et al.* 2014) in Enza-Maintained 16D$^{CRPC}$ cells with OGDHL knockdown. **(G and H)** Average expression of Gene Ontology Biological Process: Axon guidance **(G)** and Hallmark: Epithelial-mesenchymal transition **(H)** gene sets in tumors formed from 16D$^{CRPC}$ cells with genetic knockout of OGDHL (n =4) or control 16D cells (n = 5). Data represented as average Z-score. **(I and J)** Western blot of NEPC driver HES6 in control and OGDHL knockdown cells recovered 4 weeks after implantation *in vivo* **(I)** and quantification **(J)** measured as the relative band intensity normalized to Vinculin loading control.

Error bars represent +/- SEM. P-value calculated by unpaired t-test with Welch's Correction

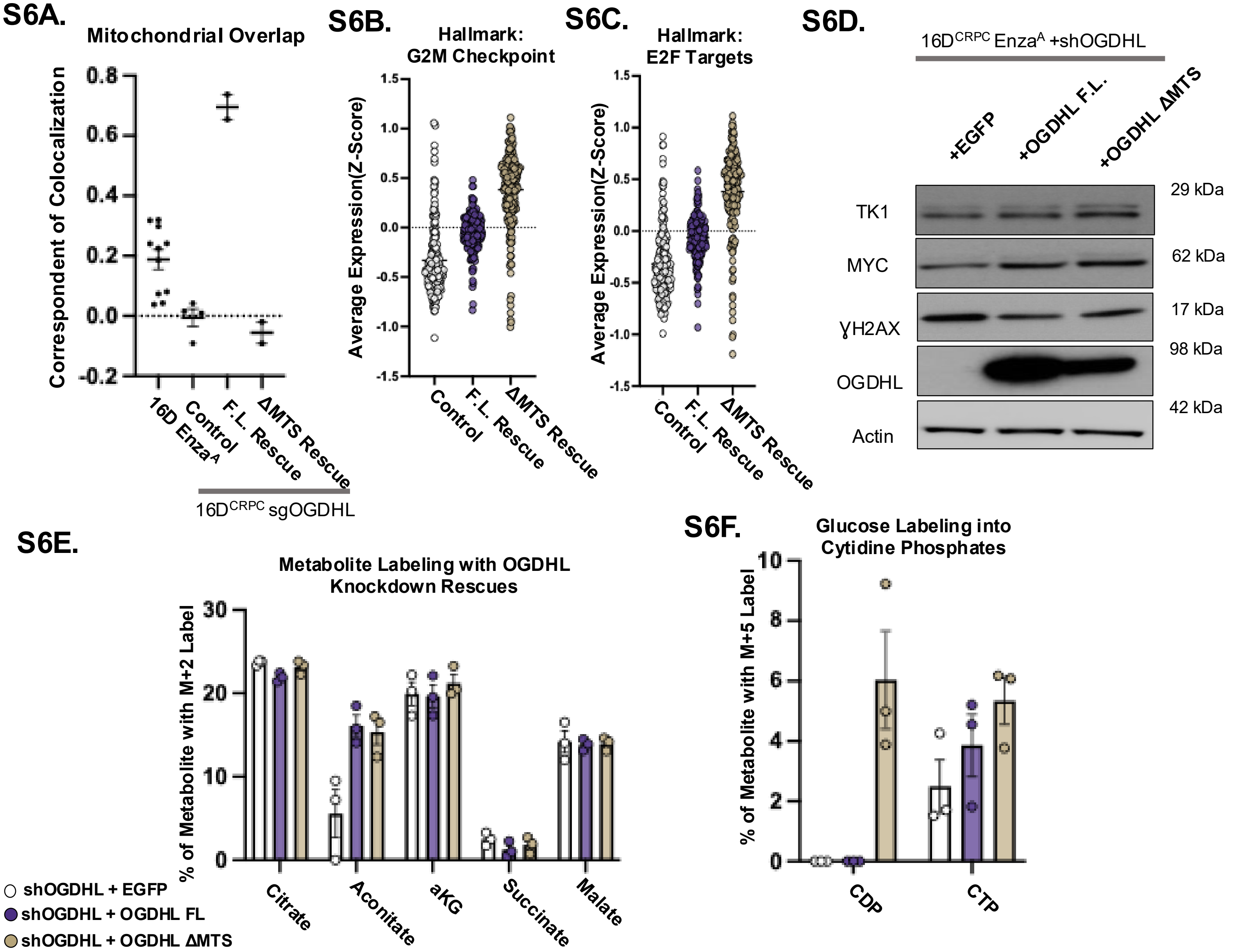


**Supplementary Data 6: Mitochondrial localization of OGDHL is not-required to regulate lineage and nucleotide phenotypes**

**(A)** Quantification of colocalization of immunofluorescence signal from OGDHL and the mitochondrial marker TUFM in Enza-maintained $16D^{CRPC}$ cells with CRISPR-Cas9 mediated genetic knockout of OGDHL and overexpression of CRISPR-resistant Full Length (FL) and mitochondrial targeting sequence deletion (ΔMTS) OGDHL variants. **(B and C)** Average expression of genes in the Hallmark: G2M Checkpoint **(B)** and Hallmark: E2F Targets gene sets **(C)** in $16D^{CRPC}$ cells with CRISPR-Cas9 mediated genetic knockout of OGDHL or reintroduction of knockout-resistant FL and ΔMTS variants. Data shown is average Z scores or TPM values from 3 technical replicates each. **(D)** Western Blot of the DNA damage marker ɣH2AX, cell cycle gene MYC and nucleotide metabolism gene TK1 in enza-maintained $16D^{CRPC}$ cells with genetic knockdown of OGDHL and expression of knockdown-resistant FL and ΔMTS variants. **(E and F)** M + 2 labeling of TCA cycle intermediates from U-$^{13}$C Glucose incorporation **(E)** and M + 5 labeling of cytidine phosphates derived from U-$^{13}$C glucose **(F)** in enza-maintained $16D^{CRPC}$ cells with genetic knockdown of OGDHL and expression of knockdown-resistant FL and ΔMTS variants. Data represents 3 technical replicates each.

Error bars represent +/- SEM. P-value calculated by unpaired t-test with Welch's Correction

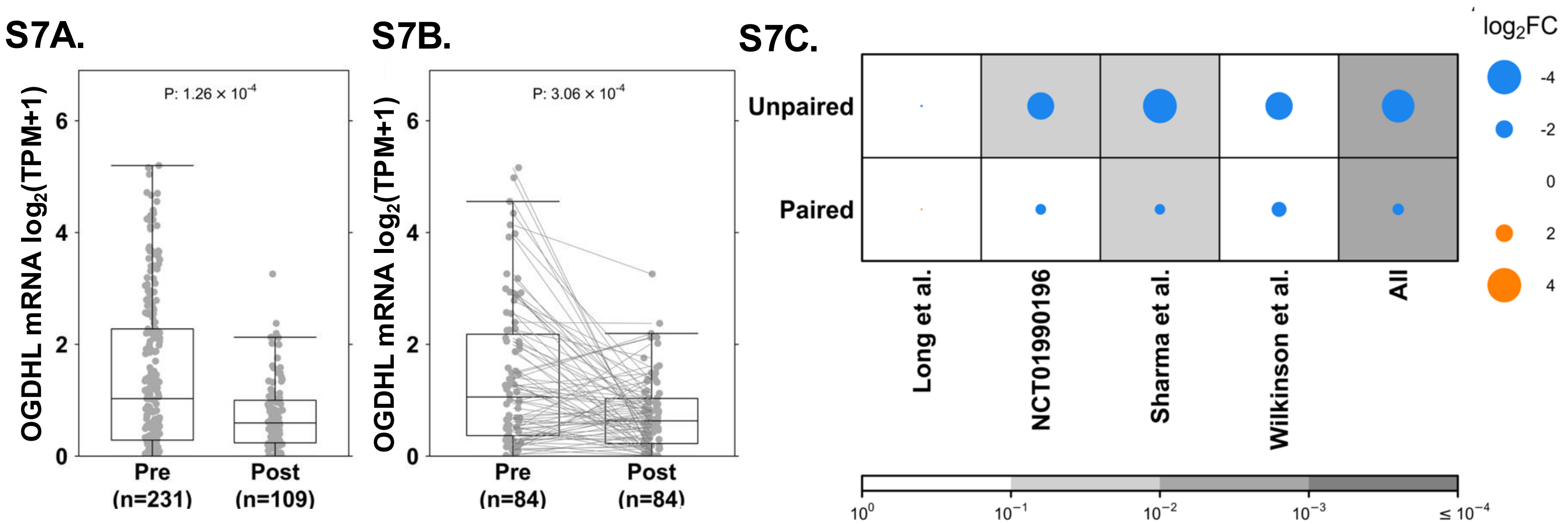


**Supplementary Data 7: Differential expression analysis of *OGDHL* across multiple studies.**
**(A)** Unpaired comparison of *OGDHL* mRNA expression levels ($\log_2$(TPM + 1)) between Pre- (n=231) and Post- (n=109) samples from aggregated datasets. Individual data points are shown as grey dots. P-value calculated using Wilcoxon rank-sum test. **(B)** Paired comparison of *OGDHL* expression levels ($\log_2$(TPM + 1)) between Pre- (n=84) and Post- (n=84) samples within the same individuals. Lines connect measurements from the same individual. P-value calculated using Wilcoxon signed-rank test. **(C)** Summary of *OGDHL* differential expression results across individual studies and combined analysis ('All'), stratified by unpaired and paired analysis types. Dot size and color indicate the $\log_2$Fold Change ($\log_2$FC) comparing Post vs. Pre conditions (key right). Background cell shading intensity reflects the statistical significance after Bonferroni multiple testing correction (scale bottom). TPM, Transcripts Per Million; *OGDHL*, Oxoglutarate Dehydrogenase L.
